# Ethambutol and Meropenem/Clavulanate Synergy Promotes Enhanced Extracellular and Intracellular Killing of *Mycobacterium tuberculosis*

**DOI:** 10.1101/2023.10.24.563807

**Authors:** Francisco Olivença, David Pires, Cátia Silveiro, Bianca Gama, Frederico Holtreman, Elsa Anes, Maria João Catalão

**Author notes:** Correspondence: Maria João Catalão and Elsa Anes.

## Abstract

Increasing evidence supports the repositioning of beta-lactams for tuberculosis (TB) therapy. However, additional research on the interaction of these drugs with conventional anti-TB agents is still warranted. Since the complex cell envelope of *Mycobacterium tuberculosis* (*Mtb*) may pose an additional obstacle to the diffusion of beta-lactams, an improved activity upon combination with drugs that inhibit the synthesis of outer cell wall elements is particularly relevant. In this context, we aimed to determine potential synergies between beta-lactams and the antimycobacterial drugs ethambutol and isoniazid. This was followed by experiments that aimed to confirm if the increased antimicrobial effects remained within the intracellular milieu and if they promoted heightened immune responses. Results of checkerboard assays with H37Rv and eight clinical isolates, including four drug-resistant *Mtb* strains, exposed that only the treatments containing ethambutol and beta-lactams achieved synergistic effects, while the standard ethambutol and isoniazid association failed to produce synergy in any of the tested isolates. In *Mtb*-infected THP-1 macrophages, combinations of ethambutol with increasing meropenem concentrations consistently displayed superior killing activities over the individual antibiotics. Flow cytometry with BODIPY FL vancomycin, which binds directly to the peptidoglycan, confirmed an increased exposure of this layer after co-treatment. This was reinforced by the high IL-1β secretion levels found in infected macrophages after incubation with concentrations of meropenem above 5 mg/L, which indicated an exposure of the host innate response sensors to pathogen-associated molecular patterns in the PG. Our findings show that the proposed impaired access of beta-lactams to periplasmic transpeptidases is counteracted by concomitant administration with ethambutol. The efficiency of this combination may be attributed to the synchronized inhibition of arabinogalactan and peptidoglycan synthesis, two key cell wall components. Given that beta-lactams exhibit a time-dependent bactericidal activity, a more effective pathogen recognition and killing prompted by this association may be highly beneficial to optimize TB regimens containing carbapenems.

## Introduction

The highly complex cell envelope of *Mycobacterium tuberculosis* (*Mtb*) is one of the most distinct structural features of the causative agent of tuberculosis (TB) and is considered essential for its antibiotic resistance and virulence (1,2). Within the envelope, the cell wall (CW) of the pathogen is organized as a layer of peptidoglycan (PG), linked to arabinogalactan (AG) polysaccharide side chains which are esterified at their distal ends with mycolic acids (MA), comprising the mycolyl-arabinogalactan-peptidoglycan (mAGP) complex (3). Unlike AG and MA, whose synthesis is inhibited by the first-line anti-TB drugs ethambutol (EMB) and isoniazid (INH), respectively, the PG sheet of *Mtb* is not usually targeted by TB therapy.

Beta-lactams prevent PG biosynthesis by blocking cross-linking enzymes such as the penicillin-binding proteins (PBPs), also known as DD-transpeptidases. However, this antibiotic class has remained excluded from routine TB therapy following the observation that *Mtb* was innately resistant to penicillin (4). This resistance has been attributed to a potentially limited beta-lactam diffusion across the lipid-rich outer layers and to the co-existence of an effective beta-lactamase, BlaC, and non-classical PG LD-transpeptidases (Ldts) (5,6). The combination with beta-lactamase inhibitors and the advent of carbapenems, a newer subclass that strongly inhibits both PBPs and Ldts, are driving the reassessment of beta-lactams as additional anti-TB agents (7,8). This is conveyed by recent clinical trials that evaluate the early bactericidal activity of beta-lactams (ClinicalTrials.gov identifiers NCT02381470, NCT02349841, and NCT03174184) and by the inclusion of two carbapenems among the drugs recommended by the WHO for use in longer multidrug-resistant TB (MDR-TB) regimens (9). While effective, the WHO considers more evidence on the role of beta-lactams within TB regimens is needed. To address this requirement, our group has explored the associations of lineage and mutational patterns of *Mtb* clinical strains with beta-lactam susceptibility and provided insights into potential drivers of resistance to these antibiotics (10,11).

TB therapeutic schemes require several antibiotics and the application of beta-lactams commands a robust understanding of how these would interact with standard drugs. Combinations of rifampicin, a cornerstone anti-TB drug, with carbapenems or faropenem were effective against clinical isolates of *Mtb* and *Mycobacterium abscessus* (12). Associations with other anti-TB drugs are still poorly defined, but EMB associated to penicillin or carbenicillin was shown to synergistically inhibit the growth of *Corynebacterium glutamicum* and *Mycobacterium phlei* (13). Like other members of the mycolata taxon, these species display a mAGP cell wall core, suggesting that beta-lactams may also interact synergistically with EMB and possibly INH against *Mtb*. This prospect is highly relevant and deserves to be investigated as these interactions may facilitate the exposure of PG and the access of beta-lactams to the transpeptidases.

Moreover, the PG is an important pathogen-associated molecular pattern (PAMP) that can be recognized by the innate immune system. We have recently shown that the *N*-glycolylation of muramic acid and the amidation of D-*iso*-glutamate, characteristic subtleties of the mycobacterial PG, contribute to beta-lactam resistance and intracellular survival of *Mycolicibacterium smegmatis* (14). PG fragments containing D-glutamyl-meso-diaminopimelic acid or muramyl dipeptide are detected by the NOD1 and NOD2 intracellular receptors, respectively (15,16). Both types of muropeptides can be found on the mycobacterial PG (17,18) and recognition by the NOD receptors leads to NF-κB-dependent upregulated expression of pro-inflammatory molecules (19). Other important pattern recognition receptors (PRRs) that recognize intracellular mycobacteria include the NOD-like receptor family, pyrin domain-containing 3 (NLRP3) and absent in melanoma 2 (AIM2). While AIM2 binds to double stranded DNA, the ligands of NLRP3 are diverse, but in both cases these innate sensors activate the irrespective inflammasomes, leading to activation of pro-IL-1β and secretion of IL-1β (20).

Beta-lactams are considered to diffuse but not to accumulate in cells because of a free carboxyl function common to all molecules (21). Thus, a possible improvement of the action of beta-lactams prompted by a higher concentration in the infection site or by a combination with other CW-targeting agents may be especially important to prevent host evasion strategies or to kill bacteria that escape the phagosome. In this work, we used a set of *Mtb* strains with diverse characteristics and anti-TB drug-resistance profiles to clarify the type of interactions established between different combinations of CW inhibitors. The main objective of this study was to decipher if beta-lactams interact synergistically with conventional antimycobacterials that also target the synthesis of CW components, i.e., EMB and INH. In addition, macrophage infection assays were employed to assess the impact of selected combinations on intracellular pathogen elimination and flow cytometry was used to verify the exposure of PG in antibiotic-treated *Mtb*. The potential influence of these antibiotic treatments on IL-1β secretion was also evaluated. The combined outputs strengthen the position of beta-lactams as alternative antimycobacterials and highlight that application of these drugs in TB therapy may greatly benefit from other CW-destabilizing agents. Moreover, they suggest that the effects of carbapenems against TB may expand outside mere antimicrobial activity against *Mtb*.

## Materials and Methods

### 1. Bacterial strains, culture conditions, and antibiotics

Eight *Mtb* clinical strains and the reference strain H37Rv were used in this study (Table S1). Antimycobacterial drug susceptibility testing under standardized guidelines, *in silico* lineage determination, and spoligotyping were previously performed for the clinical strains at the Portuguese National Institute of Health (10). Screening for bedaquiline and linezolid susceptibility was not part of standard routine during the timeframe the strains were tested. Regarding antimycobacterial drug resistance profile, clinical strains were classified according to the most recent definitions by WHO as susceptible, MDR (resistance to both rifampicin and INH), or pre-extensively drug-resistant (pre-XDR; resistance to rifampicin and any fluoroquinolone) (22). Bacteria were grown in Middlebrook 7H9 medium (BD Biosciences) or in Middlebrook 7H10 medium (BD Biosciences), supplemented with 0.2% or 0.5% of glycerol, respectively. Both media were supplemented with 10% oleic acid-albumin-dextrose-catalase (OADC) (BD Difco) and tyloxapol (Sigma-Aldrich) was added to liquid medium to a final concentration of 0.05%. Stocks of amoxicillin (AMX), EMB, MEM, and vancomycin (VAN) (Sigma-Aldrich) were prepared in purified water. Potassium clavulanate (Sigma-Aldrich) was prepared in phosphate buffer pH 6.0, 0.1 M and INH was prepared in dimethyl sulfoxide (DMSO; PanReac AppliChem).

### 2. Minimum inhibitory concentration (MIC) determination

To determine the range of concentrations to assess during the checkboard assays, the MICs of all the strains to EMB, INH, AMX/CLA, MEM/CLA were determined through an adaptation of the broth microdilution assay (23). Serial dilutions of the antibiotics were prepared in 96-well plates and bacterial cultures in the log-phase were added to a final concentration of 10^5^ CFU/mL. Wells corresponding to positive control (bacterial inoculum without any antibiotics) and negative control (without bacteria) were also included. MIC values were considered as the lowest concentration leading to no visible growth after 10-12 days of incubation of the plates at 37°C, 5% CO_2_.

### 3. Checkerboard analyses and determination of fractional inhibitory concentration (FIC) indices

Antibiotic interactions were evaluated by checkerboard assays and FIC indices were calculated to measure their extent (24). Briefly, 100 μL of supplemented liquid medium were added to the wells of a 96-well plate and the antibiotic pairs under analysis were twofold diluted along the abscissa or ordinate generating an 8 x 8 matrix. The rest of the wells were filled with medium to maintain a humidified atmosphere. 100 μL of the bacterial suspensions were added to the matrix wells to yield a final concentration of 10^5^ CFU/mL and the optical density at 600 nm (OD_600_) of each well was determined in an Infinite M200 Pro microplate reader (TECAN) after incubation at 37°C, 5% CO_2_ for 10-12 days. The MIC was considered as the concentration at which ≥99% of the growth was inhibited when compared with the positive control well. For each well that corresponded to a MIC, the FIC of each antibiotic was calculated as the MIC when the antibiotic is used in combination divided by the MIC of the antibiotic alone. The FIC index (FICI) is the sum of the individual FICs of the antibiotics under test in each well. FICI values were interpreted as follows: synergy (FICI ≤ 0.5), additive effect (0.5 < FICI ≤ 1), indifference (1 < FICI ≤ 4), and antagonism (FICI > 4) (25). We provide the minimum, median, and maximum FICI values as the mean of two independent replicates for each combination to better capture and convey the range of interactions between the tested pairs among all strains.

### 4. Growth curves and spot assays

Bacterial cultures of H37Rv or the clinical isolates at an OD_600_ of 0.5-0.6 were normalized to an OD_600_ of 0.06 by dilution with fresh 7H9 medium. Flasks were incubated for 8 days at 37°C and aliquots were collected after 1, 2, 4, 7, and 8 days of incubation to plot the growth curves.

For the growth curves in the presence of antibiotics, 100 μL of a H37Rv suspension (OD_600_ = 0.8) were added to the wells of 96-well plates containing the antibiotics to achieve an initial OD_600_ of 0.4. Wells containing each concentration of the antibiotic, the combinations or DMSO were used as respective blanks. Plates were incubated at 37°C, 5% CO_2_ and growth was monitored by measuring absorbance at 600 nm in an Infinite M200 Pro microplate reader (TECAN) at 6 h and 1, 2, 4, 6, 8, 12, and 15 days after inoculation with the bacteria. At the last timepoint, 5 μL of each suspension were spotted onto 7H10 agar plates with 10% OADC. These were incubated at 37°C and 5% CO_2_ until spots were visible.

### 5. Macrophage assays

#### 5.1. Cell culture and maintenance

Human promonocytic THP-1 cells were maintained at an initial density of 5 × 10^5^ cells/mL in RPMI 1640 medium (Cytiva) supplemented with 10% fetal bovine serum (Corning) in a humidified atmosphere at 37°C and 5% CO_2_. When required, 5 × 10^5^ cells/mL were seeded in the wells of flat-bottom tissue culture plates and matured into macrophages with 20 nM phorbol 12-myristate 13-acetate (PMA; Sigma-Aldrich) during 72 h. Prior to any assay with the differentiated macrophages, medium with PMA was removed and replenished with the appropriate prewarmed cell culture medium for each assay. The 96-well format (5 × 10^4^ cells/well) was used for CFU enumeration and resazurin viability assays, while the 24-well format (2.5 × 10^5^ cells/well) was selected for cell death assays.

#### 5.2. Infection with *Mtb*

To assess the intracellular efficacy of the drugs, cultures of the reference and the clinical strains on the exponential growth phase were centrifuged, washed, and resuspended in macrophage culture medium. These suspensions were subjected to an ultrasonic bath for 5 min to dismantle aggregates and residual clumps were removed by centrifugation at 500 *g* for 1 min (26). Final single-cell bacterial suspensions were prepared and macrophages were infected at a multiplicity of infection of 1 for 3 h at 37°C with 5% CO_2_. Following internalization, culture medium containing bacteria was discarded and remaining extracellular bacteria were removed by gently and thoroughly washing the cells with prewarmed phosphate-buffered saline (PBS). Cells were then overlaid with prewarmed culture medium, with or without antibiotics.

To assess IL-1β secretion in the different conditions of drug treatment in infected macrophages, a single-cell suspension of *Mtb* H37Rv was adjusted to a final concentration equal to the one used during the internalization step of previous infection assays. Aliquots of 1 mL were treated with individual antibiotics (EMB at 2 mg/L; MEM/CLA at 0.5 mg/L) or in combination. When present, CLA concentration was always fixed at 2.5 mg/L. Following 6 h of incubation at 37°C with 5% CO_2_, untreated and treated aliquots were centrifuged and washed twice with 1 mL of fresh culture medium. These suspensions were then used to infect THP-1 macrophages differentiated in 96-well plates as described above. Prewarmed culture medium without antibiotics was added to the macrophages after internalization.

#### 5.3. Intracellular survival evaluation by CFU quantification assays

Estimation of intracellular live bacilli after drug treatment was performed as previously described (26,27). Briefly, infected macrophages were lysed for 15 min with 0.05% IGEPAL solution (Merck) after 3 h of internalization (day 0) and after 1, 3, and 5 days of infection. The lysates were serially diluted and plated onto supplemented 7H10 plates and incubated at 37°C with 5% CO_2_ for two to three weeks until colonies could be observed and enumerated under the microscope. The average growth of strains in defined groups after the different treatments was plotted over time as the logarithm of the ratio between the CFU/mL in each timepoint over the CFU/mL on day 0. Within individual timepoints, average CFU/mL yielded by each treatment was compared with the untreated control.

### 6. Apoptosis and necrosis assay using flow cytometry

For identification of apoptotic and necrotic cells after 72 h of antibiotic treatment, FITC Annexin V Apoptosis Detection and Zombie Red Fixable Viability kits (Biolegend) were used following the instructions of the manufacturer. Briefly, the supernatants of macrophages infected with *Mtb* H37Rv or strain MDR III were collected. Uninfected macrophages, either untreated or treated with DMSO at 1% (v/v), were included as controls. Cells were detached with accutase (Corning) for 15 min and then pooled with the respective supernatants. Cells were centrifuged at 500 *g* for 5 min at room temperature, and the pellets were resuspended and incubated with FITC Annexin V and Zombie Red in the dark for 15 min. Cells were washed with the appropriate kit buffer and fixed in 4% paraformaldehyde (PFA; Thermo Scientific) in PBS for 1h. Following fixation, cells were again washed and resuspended with the buffer. Samples were analyzed by flow cytometry using a Cytek Aurora flow cytometer (Cytek Biosciences) and data analysis was conducted in FCS Express 7 (De Novo Software).

### 7. Viability assays

Macrophages seeded in 96-well plates at a concentration of 5 × 10^5^ cells/mL (100 μL per well) were treated with different antibiotic combinations for 1, 3, or 5 days. At each timepoint, cell medium was removed and prewarmed fresh medium with 10% resazurin (TCI Chemicals) was added. After 3 h of incubation at 37°C and 5% CO_2_, fluorescence emission was analyzed in an Infinite M200 Pro microplate reader (TECAN) according to the instructions of the manufacturer. Non-treated macrophages were used as reference and cells treated with DMSO at 1% (v/v) were used as control for cell death.

### 8. Evaluation of peptidoglycan exposure with BODIPY FL vancomycin

Cultures of *Mtb* H37Rv and of the clinical strains Susceptible III and MDR III on the exponential growth phase were diluted to an OD_600_ of 0.1 with supplemented 7H9 medium. The bacterial suspensions were treated with individual antibiotics (EMB at 2 mg/L; MEM/CLA at 0.5 mg/L) or in combination. Strain MDR III was additionally treated with EMB at 16 mg/L, either individually or in association with MEM/CLA. After treatment for 6 h at 37°C with 5% CO_2_, 1 mL aliquots were collected, centrifuged at 3000 *g* for 5 min, and resuspended in PBS 1x with 1 mg/L BODIPY FL vancomycin (Invitrogen) (28) and 0.5 µM BacLight Red (Invitrogen). Susceptibility to VAN was previously determined to confirm that 1 mg/L of BODIPY FL vancomycin was below the MIC in the three strains. Staining was performed for 15 min at 37°C in the dark, followed by centrifugation. Pellets were washed twice with PBS 1x to remove remaining free dye and then fixed with 4% PFA in PBS 1x for 30 min. The samples were centrifuged, resuspended in PBS 1x, and analyzed by flow cytometry using a Cytek Aurora flow cytometer (Cytek Biosciences). Data analysis was performed in FCS Express 7 (De Novo Software). Fixed samples of *Mtb* H37Rv were mounted with ProLong™ Gold antifade mountant (Invitrogen) and analyzed by confocal microscopy (Leica AOBS SP5).

### 9. IL-1β Quantification

The supernatants of the macrophage cultures infected with strains *Mtb* H37Rv, susceptible III, and MDR III were collected after 24 h of treatment with the antibiotics and stored at −80°C for posterior evaluation of IL-1β secretion by Sandwich Enzyme-Linked Immunosorbent Assay (ELISA). In addition, the supernatants retrieved after 24 h of infection with *Mtb* H37Rv pre-treated with selected antibiotic concentrations were also tested. The supernatants collected from uninfected macrophages, either untreated or incubated with antibiotics or lipopolysaccharide (LPS; Sigma-Aldrich) at 100 ng/mL for 24 h, were added as controls. Quantification was conducted using ELISA Max Deluxe Set Human IL-1β kit (Biolegend) according to the instructions of the manufacturer. Absorbance was measured at 570nm and 450nm in a Varioskan LUX multimode microplate reader (Thermo Scientific).

### 10. Statistical analysis

Heatmaps and statistical analysis, including one-way ANOVA for multiple group comparisons, were performed in GraphPad Prism version 9.0.

## Results

### 1. The selected set of strains displayed diverse antibiotic susceptibility and growth phenotypes

Since susceptibility to beta-lactams is not currently part of routine testing, we have determined the MICs of EMB, INH, AMX/CLA, and MEM/CLA by an adaptation of the broth microdilution assay (23) to obtain these values under the same conditions for all strains (Table S1). The results were fully in line with previous resistance classification of the strains by Bactec MGIT960 (Becton Dickinson), with four resistant isolates (MDR I-III and pre-XDR) having MICs of the anti-TB drugs above the standard concentrations used in this method (INH - 0.1 mg/L; EMB - 5.0 mg/L). Strains Susceptible II and MDR III had beta-lactam MICs higher than H37Rv, while strains MDR I, Susceptible III, Susceptible IV, and pre-XDR had lower AMX/CLA MICs. Strains MDR I and Susceptible III additionally had an MIC of 0.25 mg/L of MEM/CLA.

Seven strains belonged to lineage 4 (sublineages 4.1.2.1 (*n*=1), 4.2.1 (*n*=2), and 4.3.4.2 (*n*=4)) and the remaining strain was from lineage 2 (sublineage 2.2.1.1). The extracellular and intracellular growth of the different stains was then assessed. In 7H9 broth medium, apart from strain MDR I, all strains from sublineages 4.2.1 and 4.3.4.2 presented a slower growth than the reference strain from day 4 onward (Figure S1A). Susceptible strains I and II grew faster than the other strains throughout the entire assay. Compatible with what was observed extracellularly, isolates from sublineage 4.3.4.2 mostly displayed a lower intracellular growth than strains from sublineages 2.2.1.1 and 4.1.2.1 (Figure S1B), except for strain Susceptible IV. However, not all patterns were mirrored, with sublineage 4.2.1 strains undergoing more pronounced increases in their relative intracellular growth than sublineage 4.3.4.2 strains. The number of strains is not enough to formulate broader assumptions on possible correlations between sublineage and growth pattern, but our results demonstrate that the group of strains included in the present study showed different combinations of anti-TB/beta-lactam susceptibility and growth profiles.

### 2. Ethambutol and beta-lactams interact synergistically

Synergy assays combining the two anti-TB drugs, the two beta-lactams, or one anti-TB drug and one beta-lactam were then performed. The relative growth at the end of the assay was assessed (Figure 1) and the FICI was calculated for wells with less than 1% growth of the control (Figure S2). The lowest FICI (FICI_Min_), the median FICI (FICI_Med_), and the highest FICI (FICI_Max_) were calculated for each combination and in each strain (Figures 2; S3). Additive effects were noted for all associations, but synergies were only detected for EMB combined with AMX/CLA or MEM/CLA. For each of these combinations, 5/9 strains had a FICI_Min_ < 0.5 and two strains also presented a FICI_Med_ compatible with synergy. An interesting pattern was noticed for EMB plus MEM/CLA, with all strains with FICI_Min_ < 0.5 belonging to sublineage 4.3.4.2. One of these isolates was the pre-XDR strain in our study, which tended to have lower FICI values. Nonetheless, with values for FICI_Min_ and FICI_Med_ < 0.5 and FICI_Max_ < 1, this strain seemed to be particularly vulnerable to the EMB and beta-lactam combinations. Despite not showing synergy, strains Susceptible I-II and MDR III still presented the lowest FICI_Min_ values (< 0.7) for the associations between EMB and a beta-lactam antibiotic.

**Figure 1.**
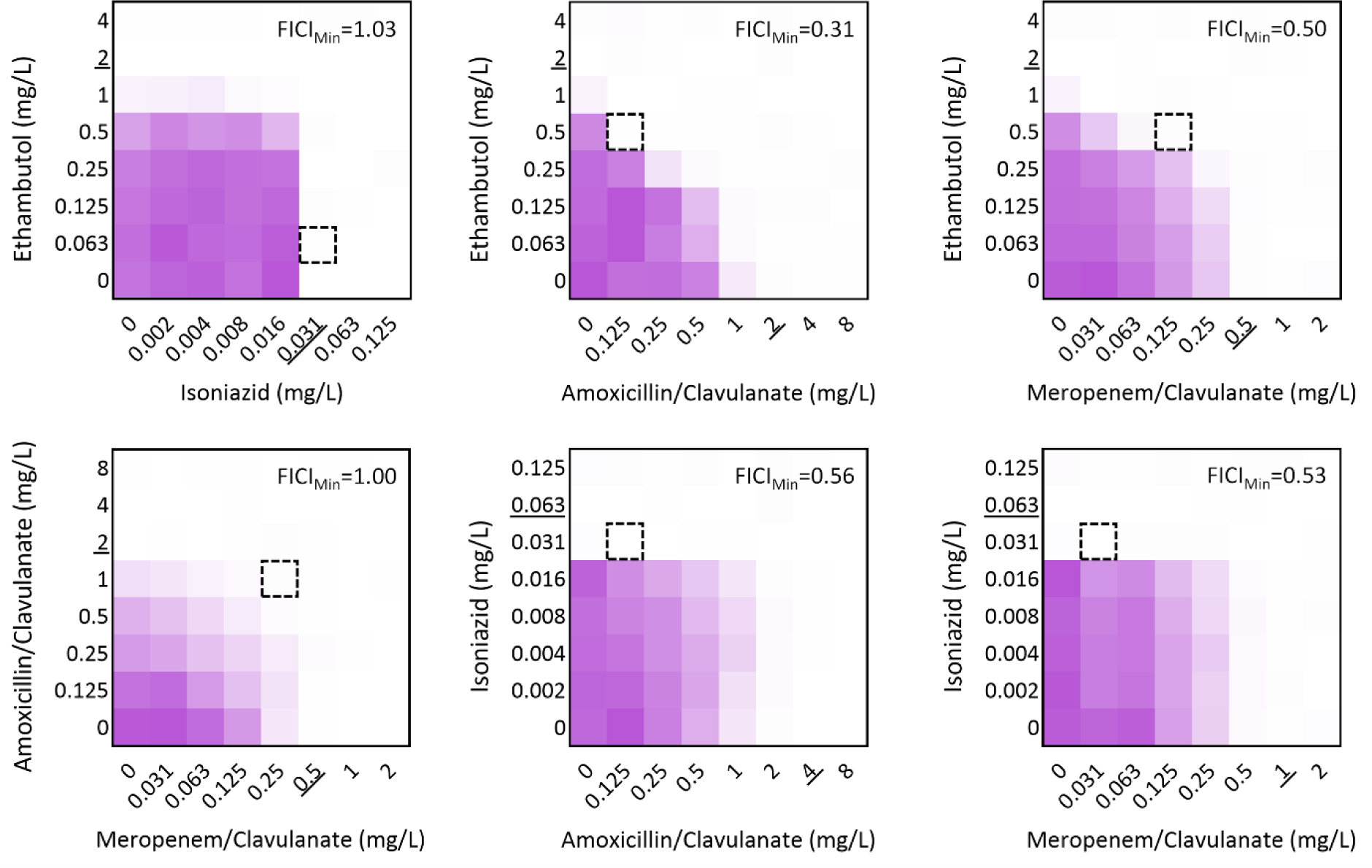
Heatmap depiction of the outputs of the checkerboard assays in *Mycobacterium tuberculosis* H37Rv. Data represents the optical density at 600 nm (OD_600_) of each well compared to the control for one representative biological replicate. Dark purple regions represent higher cell density. Underlined values show the minimum inhibitory concentration of the antibiotic in the respective assay. The dashed square indicates the combination that yielded the lowest FIC index value (FICI_Min_). When present, clavulanate concentration was fixed at 2.5 mg/L.

**Figure 2.**
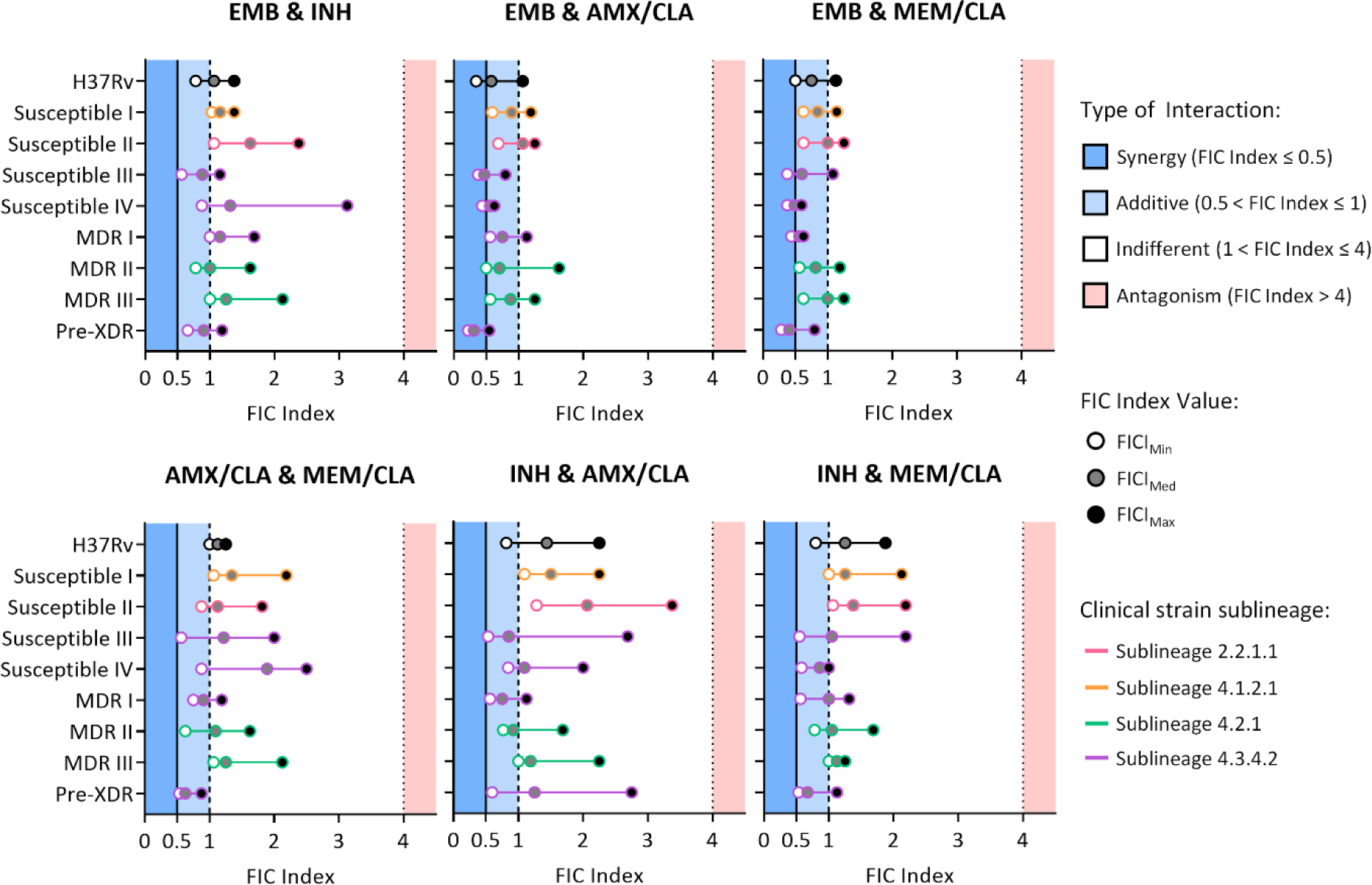
Representation of the FIC index values calculated from the interactions established between different antibiotics in *Mycobacterium tuberculosis* H37Rv and eight clinical isolates. Dots represent the lowest FICI (FICI_Min_), the median FICI (FICI_Med_), and the highest FICI (FICI_Max_) calculated as the mean of two independent replicates for each combination and in each strain. The color of the outline of the dots denotes the respective sublineage of the clinical strains. AMX, amoxicillin; CLA, clavulanate; EMB, ethambutol; INH, isoniazid; MEM, meropenem.

Importantly, the addition of one beta-lactam to EMB exceeded the performance of the standard therapeutic association of EMB with INH, which failed to produce any synergy, even in fully susceptible strains. For two isolates (Susceptible IV and MDR I) this combination actually resulted in the highest FICI_Max_. Similarly, INH combined with a beta-lactam or the dual beta-lactam treatment did not yield synergistic effects. The FICI_Min_ obtained for INH with either AMX/CLA or MEM/CLA was generally lower or on par with the attained for INH combined with EMB. The FICI_Max_ was sometimes higher for INH with the beta-lactams but not high enough to induce antagonism.

Overall, no antagonism effect was observed for any drug combination. In addition, beta-lactams do not appear to hinder the effect of the anti-TB drugs and interact synergistically with EMB in some strains. To select the most promising EMB/beta-lactam combination for the following *in vivo* infection assays, we took into consideration the individual properties of the two beta-lactams. Amoxicillin, either combined or not with clavulanate, requires higher concentrations to kill *Mtb* than meropenem (10,23,29). In addition, this carbapenem is also more resilient to hydrolysis by BlaC (5,30). In our studies, INH combined with MEM/CLA also had generally lower FICI values than INH plus AMX/CLA. Notably, of the two beta-lactams, meropenem is the only one currently included by WHO for longer regimens for DR-TB. For these reasons, EMB with MEM/CLA was the preferred combination for further studies.

### 3. Co-treatment with ethambutol and meropenem results in a faster and more sustained intracellular pathogen clearance

Infection assays were conducted to verify if the synergy between EMB and MEM/CLA was maintained intracellularly. Cells were infected with the nine strains and EMB and/or MEM/CLA were added after the internalization period. For meropenem, multiples of the MIC for H37Rv were tested (0.5, 5, and 50 mg/L) and clavulanate was added to a final concentration of 2.5 mg/L. For all strains, EMB was tested at 2 mg/L, the MIC for H37Rv. A concentration of 16 mg/L was additionally assessed with the resistant strains, as this was a median value for the four EMB-resistant isolates. This way, three groups were defined: EMB-susceptible (EMB*^S^*) strains treated with 2 mg/L EMB; EMB-resistant (EMB*^R^*) strains treated with 2 mg/L EMB; EMB*^R^*strains treated with 16 mg/L EMB.

The kinetics of the infection with the EMB*^S^* strains with 2 mg/L EMB (Figure 3A) revealed that the lowest MEM/CLA concentration had a marginal effect compared to the control and that on day 5 the growth had decreased from the previous timepoint in both conditions, possibly due to macrophage functional impairment. While the addition of 5 mg/L of MEM/CLA stabilized growth until day 3, 50 mg/L of the beta-lactam at this timepoint accomplished a reduction in growth close to 10% of the initial value. Individually, EMB seemed to have a slightly slower start of action than the highest MEM/CLA concentration but over time EMB steadily decreased growth and attained almost the same endpoint value as this treatment. All the combined treatments resulted in a lower growth than the respective individual antibiotics, with EMB combined with 5 mg/L and 50 mg/L of MEM/CLA having the steepest decreases and achieving a relative growth on day 5 of only 5% and 1.5%, respectively.

**Figure 3.**
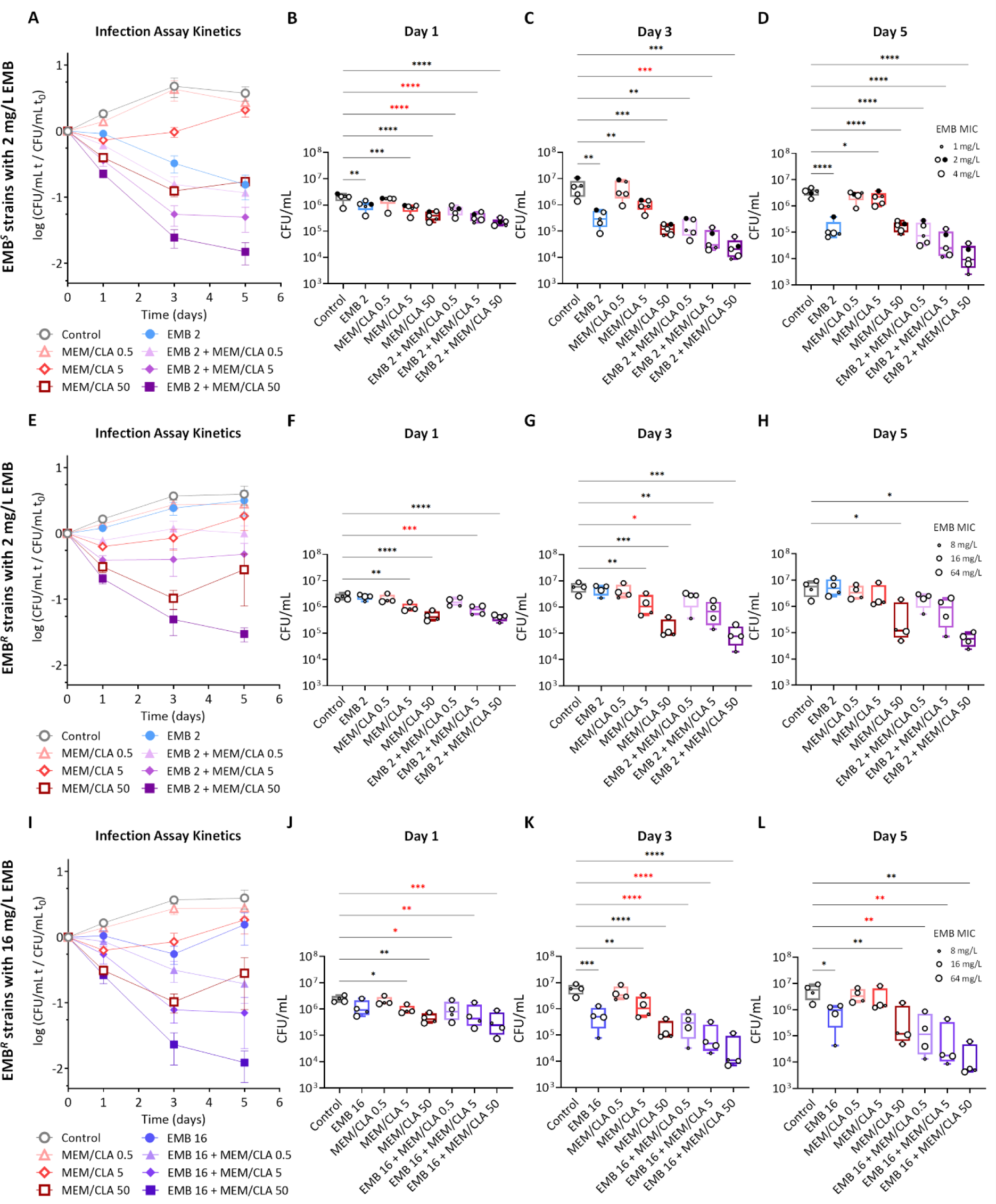
Intracellular growth of *Mycobacterium tuberculosis* (*Mtb*) H37Rv and eight clinical strains in a THP-1 infection model. (A) (E) (I) Kinetics of the mean intracellular growth of ethambutol-susceptible (EMB*^s^*) (*n*=5) or ethambutol-resistant (EMB*^R^*) (*n*=4) strains with 2 mg/L or 16 mg/L of ethambutol (EMB) over 5 days of infection. The log of the ratio between the CFU/mL in each timepoint over the CFU/mL at day 0 is plotted. Results represent the average of all strains, each with at least 3 replicates per timepoint. Error bars show the standard error of the mean. (B-D) (F-H) (J-L) Intracellular bacterial counts for each strain group at day 1, 3, and 5 of the infection assay. Boxplots represent the group mean values and error bars the minimum and maximum values. Dots show absolute enumeration for individual strains (black dots (●) - *Mtb* H37Rv; white dots (○) - clinical strains), each with at least 3 replicates per timepoint. The size of the dots represents the minimum inhibitory concentration (MIC) of EMB of each strain. Statistical differences were analysed by an ANOVA test: * when *p* ≤ 0.05; ** when *p* ≤ 0.01; *** when *p* ≤ 0.001; **** when *p* ≤ 0.0001. Combined treatments that had more statistically significant effects than both unconjugated antibiotics are labeled in red. CLA, clavulanate; MEM, meropenem. The numbers after each antibiotic abbreviature indicate the respective concentration in mg/L. When present, clavulanate concentration was fixed at 2.5 mg/L.

During infection with the EMB*^R^* strains, the profile of the curves following incubation with the three concentrations of MEM/CLA mimics the obtained for the susceptible strains (Figure 3E). Contrarily, treatment with 2 mg/L of EMB did not have any impact in the infection kinetics. This absence was expected and seemed to severely affect the extent of the combinations with the two lowest MEM/CLA concentrations on later timepoints. When treated with 16 mg/L of EMB, growth was inhibited until day 3, but EMB*^R^* strains still managed to overcome the inhibitory effect at the last timepoint (Figure 3I). However, increasing the concentration of EMB shifted the kinetic profiles of all the combinations with the carbapenem to a level in which they essentially overlap the ones determined for the EMB*^S^*strains. Strain MDR III persistently displayed the highest CFU/mL counts. On the other hand, strain MDR I, which had the lowest EMB MIC (8 mg/L) among the EMB*^R^* set, tended to yield the lowest counts in co-treatments with any of the EMB concentrations.

Within each timepoint, the combined treatments that had a more statistically significant effect than both unconjugated antibiotics (labeled in red in Figure 3B-D, F-H, J-L) were EMB (either at 2 or 16 mg/L) with 0.5 mg/L or 5 mg/L of MEM/CLA, and occurred mostly on days 1 and 3. An association with 50 mg/L of MEM/CLA only presented a more significant effect compared with its individual components in the EMB*^R^* isolates treated with 16 mg/L of EMB on day 1 post-infection (Figure 3J). On day 5, the effects of the combinations never displayed a greater statistical significance, except for 16 mg/L of EMB with the two lowest MEM/CLA concentrations (Figure 3L).

Collectively, these assays suggest that the synergy between EMB and MEM/CLA is at least partially maintained in the macrophage infection model, leading to faster bacilli clearance, and suggest that resistance to EMB is an important factor for the intracellular effect of this interaction.

### 4. THP-1 viability and programmed cell death activation are unaffected by the antibiotics

After observing that the antibiotics synergized in the infection model, we wanted to understand if the tested concentrations had any cytotoxic effects in the THP-1 cells. We first analyzed uninfected macrophages treated with different concentrations of EMB and MEM/CLA, both individually and in combination, at timepoints that mimicked the infection assay. The results of the resazurin assays showed that over the 5 days of incubation, viability after the antibiotic treatments tended to slightly decrease, but never surpassing a 10% reduction (Figure 4A). In each specific timepoint, increasing concentrations of meropenem did not have any impact in cell viability, but cells in the presence of 2 mg/L of EMB had a higher fluorescence signal than the control or 16 mg/L on days 3 and 5.

**Figure 4.**
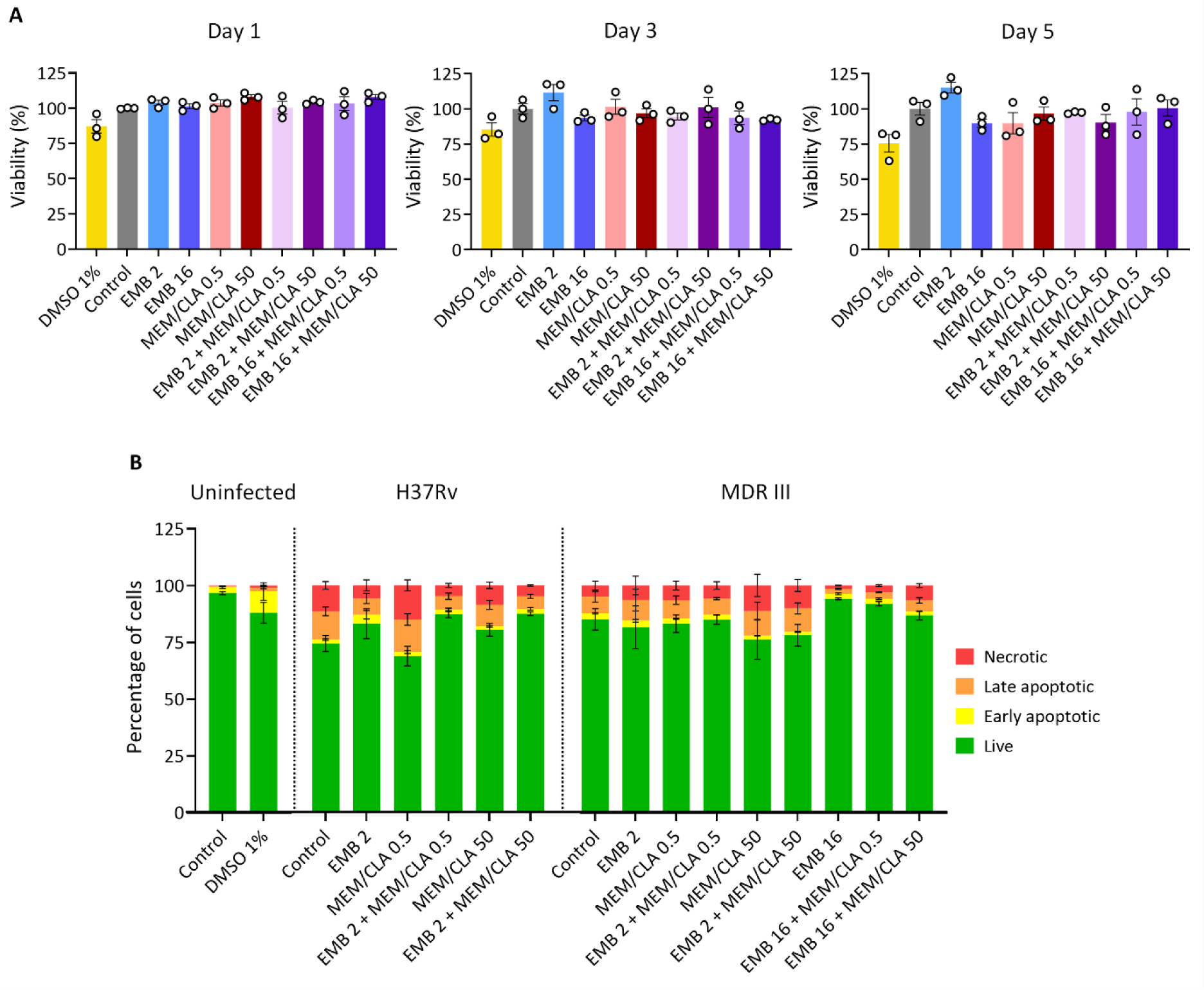
Effect of the individual or combined antibiotic treatments on the viability and programmed cell death of THP-1 macrophages. (A) Viability of uninfected THP-1 macrophages was evaluated over 5 days by measuring fluorescence intensity after the addition of resazurin. Results were calculated relative to the control (untreated macrophages - 100% viability). Bars represent the mean of three independent experiments and the error bars depict the standard error of the mean. (B) Flow cytometry analysis of the percentage of uninfected or infected macrophages stained for FITC Annexin V and/or Zombie Red at day 3 post-infection. Results represent the mean of three biological replicates and error bars show the standard error of the mean. CLA, clavulanate; EMB, ethambutol; MEM, meropenem. The numbers after each antibiotic abbreviature indicate the respective concentration in mg/L. When present, clavulanate concentration was fixed at 2.5 mg/L.

Next, we resorted to flow cytometry to assess the impact of selected treatments on programmed cell death in *Mtb*-infected macrophages after 3 days of incubation (Figures 4B; S4; S5; S6). FITC Annexin V and Zombie Red were used as markers for apoptosis and necrosis, respectively. In cells infected with the reference strain H37Rv, apart from MEM/CLA at 0.5 mg/L, all treatments increased the percentage of live cells and reduced the late apoptotic and necrotic populations. In the infection with strain MDR III, 16 mg/L EMB seemed to have a protective effect on the macrophages, increasing live cells by almost 10%. On the other hand, the highest concentration of MEM/CLA increased necrosis. This opposing tendency was equally noticed for the association of this treatment with the higher concentration of EMB. Nonetheless, both the resazurin and the flow cytometry experiments indicate that the different treatments did not have a substantial impact on cell viability or programed cell death activation that could confound the interpretation of the outputs of the infection assays.

### 5. Meropenem-induced binding of BODIPY FL vancomycin to peptidoglycan is enhanced by ethambutol

It has been previously shown that exposure of *Mtb* to meropenem increases the binding of BODIPY FL vancomycin (28). Given that we detected a synergy between EMB and MEM/CLA, and that both antibiotics inhibit the synthesis of key mycobacterial cell wall components, we aimed to verify if the association could enhance the effects of the carbapenem due to improved access. To achieve this, we treated three strains (H37Rv, Susceptible III, and MDR III) with antibiotics, stained them with BODIPY FL vancomycin, and processed the samples through flow cytometry. The BODIPY FL vancomycin concentration used was below the MIC of vancomycin of all strains (Table S1).

As expected, incubation with 0.5 mg/L MEM/CLA significantly increased BODIPY FL vancomycin fluorescence in all strains (Figures 5A-B; S7; S8; S9). This phenomenon was proportional to the basal fluorescence level of each untreated strain, with the reference and Susceptible III strains achieving the highest and lowest values, respectively. In our study, EMB alone only generated a significant effect in strain Susceptible III, but the addition of this antibiotic to MEM/CLA resulted in fluorescence increases in the three strains, albeit only statistically significant for the clinical strains. Exposure of strain MDR III to treatments with 16 mg/L EMB did not augment this effect and essentially resulted in the same fluorescence intensities as the lowest concentration. Confocal microscopy with *Mtb* H37Rv as a representative strain was consistent with the flow cytometry outputs and revealed increased BODIPY FL fluorescence following treatments with MEM/CLA, mostly concentrated at the poles (Figure 5C).

**Figure 5.**
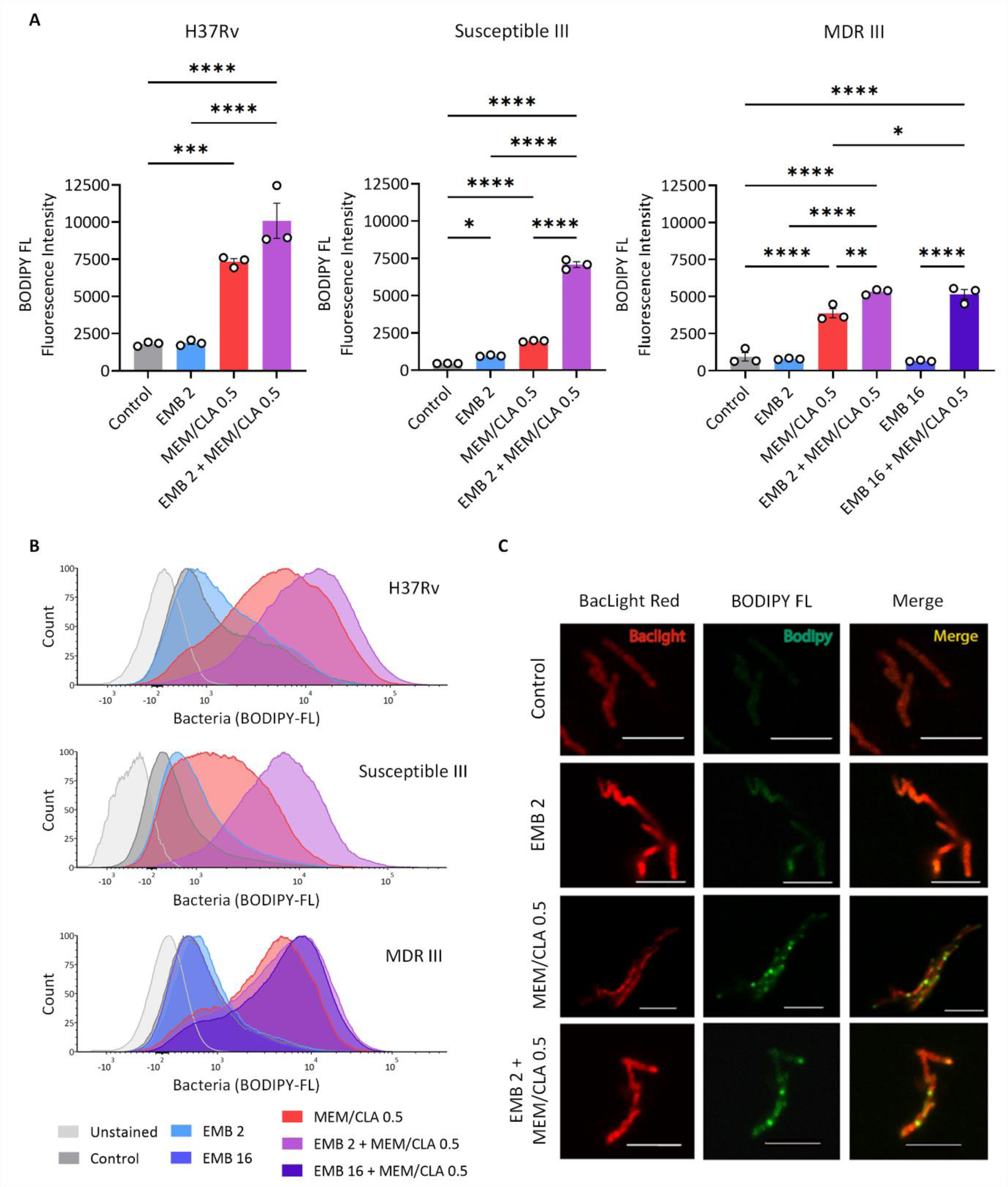
Binding of BODIPY FL vancomycin to *Mycobacterium tuberculosis* (*Mtb*) peptidoglycan. (A) Flow cytometry analysis of three *Mtb* strains (H37Rv, Susceptible III, and MDR III) untreated (control) or treated with 2 mg/L ethambutol (EMB), 0.5 mg/L meropenem/clavulanate (MEM/CLA), or both antibiotics after 6h. For MDR strain III the concentration of 16 mg/L EMB and the respective combination with MEM/CLA were also tested. The bars represent the mean BODIPY FL fluorescence intensity and the error bars depict the standard error of the mean calculated from three independent experiments. (B) Histograms display one representative experiment for each strain. (C) Fluorescence microscopy panel for *Mtb* H37Rv. Scale bar: 5 µm. The numbers after each antibiotic abbreviature indicate the respective concentration in mg/L. When present, clavulanate concentration was fixed at 2.5 mg/L.

Our findings suggest that the presence of EMB, even if at a lower concentration, promotes the destabilization of the cell wall of *Mtb* strains and contributes to nascent PG exposure, which further enhances the effects of meropenem against this pathogen.

### 6. Increasing the concentration of meropenem impacts IL-1β secretion by *Mtb*-infected macrophages

Since the antibiotic treatments compromise the integrity of the bacterial cell wall and potentially increase PG exposure, we investigated if they could also impact host recognition by measuring IL-1β secretion levels through ELISA. First, we incubated uninfected macrophages under different conditions and confirmed that the antibiotics on their own did not cause any relevant changes to IL-1β secretion (Figure S10). Next, using infection assays with the same set of strains from the previous section, we found that secretion upon 24 h of incubation with EMB and increasing MEM/CLA concentrations just mirrored the effects of the various MEM/CLA conditions applied individually (Figure 6). In some cases, combining the two drugs actually yielded a lower cytokine secretion. The most notorious of these situations occurred for the association of 16 mg/L EMB with 5 mg/L MEM/CLA in the infection with strain MDR III. These observations suggest that the combined treatment may not significantly affect host recognition.

**Figure 6.**
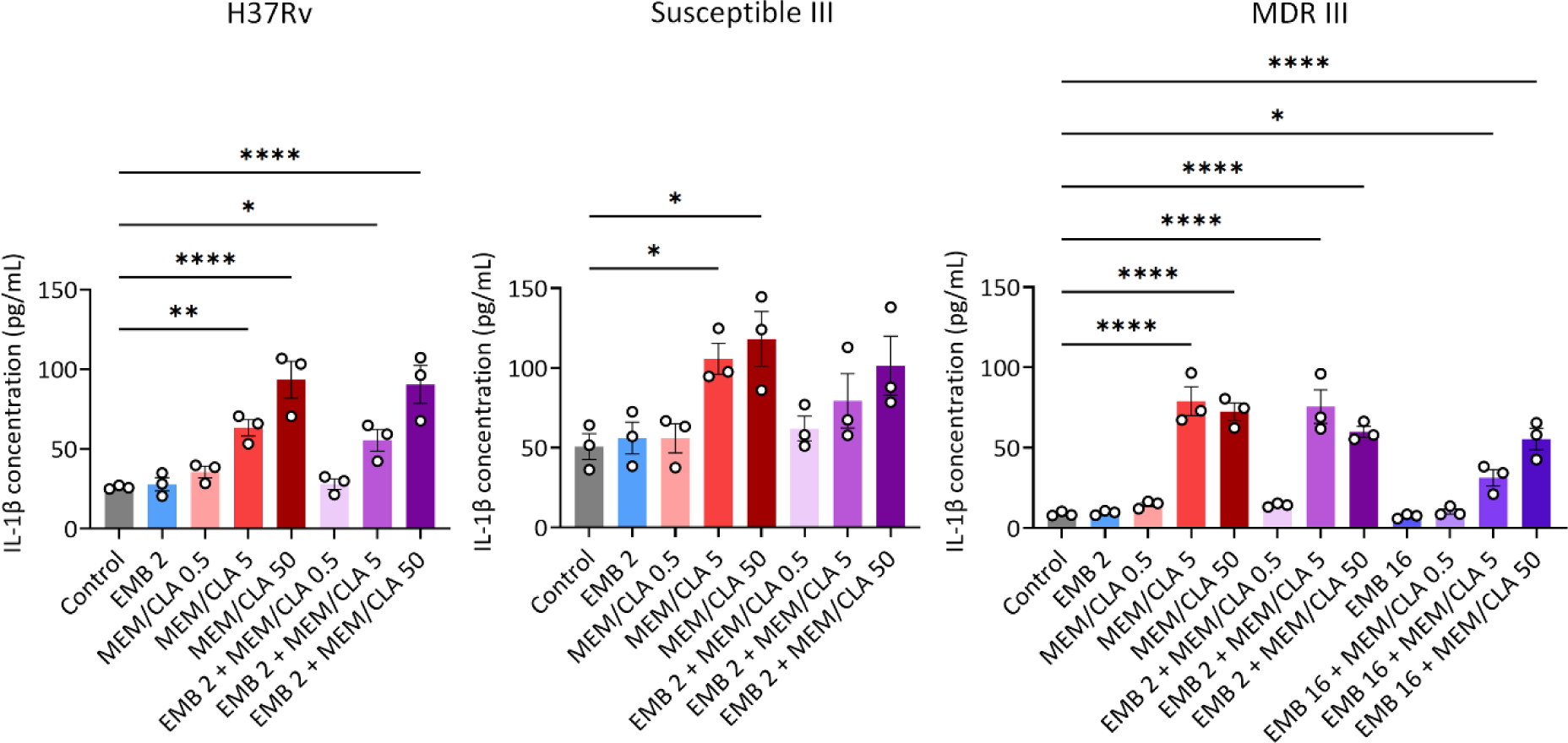
IL-1β quantification by ELISA in the supernatants of THP-1 macrophages infected with three strains (H37Rv, Susceptible III, and MDR III) after 24h of incubation with the antibiotics. Depicted values show the mean concentration of three replicates and error bars represent the standard error of the mean. Statistical differences were analysed by an ANOVA test: * when *p* ≤ 0.05; ** when *p* ≤ 0.01; *** when *p* ≤ 0.001; **** when *p* ≤ 0.0001. CLA, clavulanate; EMB, ethambutol; MEM, meropenem. The numbers after each antibiotic abbreviature indicate the respective concentration in mg/L. When present, clavulanate concentration was fixed at 2.5 mg/L.

On the other hand, we consistently observed significant increases in IL-1β secretion after incubation with 5 or 50 mg/L MEM/CLA, even in macrophages infected with strain MDR III, which had a particularly low basal cytokine secretion. We performed an additional ELISA on supernatants obtained 24 h after infection with *Mtb* H37Rv pre-treated with the lower concentrations of EMB and MEM/CLA, individually or combined. While the levels of secreted IL-1β were similar to the ones obtained with the respective treatments applied after the internalization step, significant statistical differences versus the control were observed for treatments containing the lowest MEM/CLA concentration (Figure S11). This is compatible with the outputs of the experiments with BODIPY FL vancomycin and suggests that a minimal concentration of meropenem is required to induce alterations in the IL-1β secretion levels when the carbapenem is applied post-infection. Once again, the addition of EMB to MEM/CLA did not significantly enhance this effect. Thus, meropenem concentration emerges as the main driver of IL-1β induction.

### 7. Ethambutol effectively prolongs the effects of meropenem

After observing that MEM/CLA increases IL-1β secretion by *Mtb*-infected THP-1 macrophages, we sought to observe if there were differences in the pathogen growth in the presence of the antibiotics that could suggest lysis and if these effects varied with the concentration of the antibiotics or duration of exposure. We assessed the growth of *Mtb* H37Rv by optical density measurements (Figure 7). DMSO 20% (v/v) was used as a control, as concentrations of the organic solvent above 10% have been shown to cause severe growth inhibition of *Mtb* and other bacteria (31,32). In the presence of increasing EMB concentrations, density continued increasing, but at day 8 the growth started to lag, especially with concentrations equal to or above the MIC (Figure 7A). When treated with MEM/CLA, while all conditions apart from 0.125 mg/L decreased density in the first two days, only suprainhibitory concentrations promoted a sustained decline compatible with bacterial lysis (Figure 7B). Our results are consistent with observations by Kumar et al. (33), that report *Mtb* lysis upon treatment with MEM/CLA. The more modest effects caused by 0.125 or 0.5 mg/L of MEM/CLA may result from the fact that we used cultures at a higher initial density than these authors.

**Figure 7.**
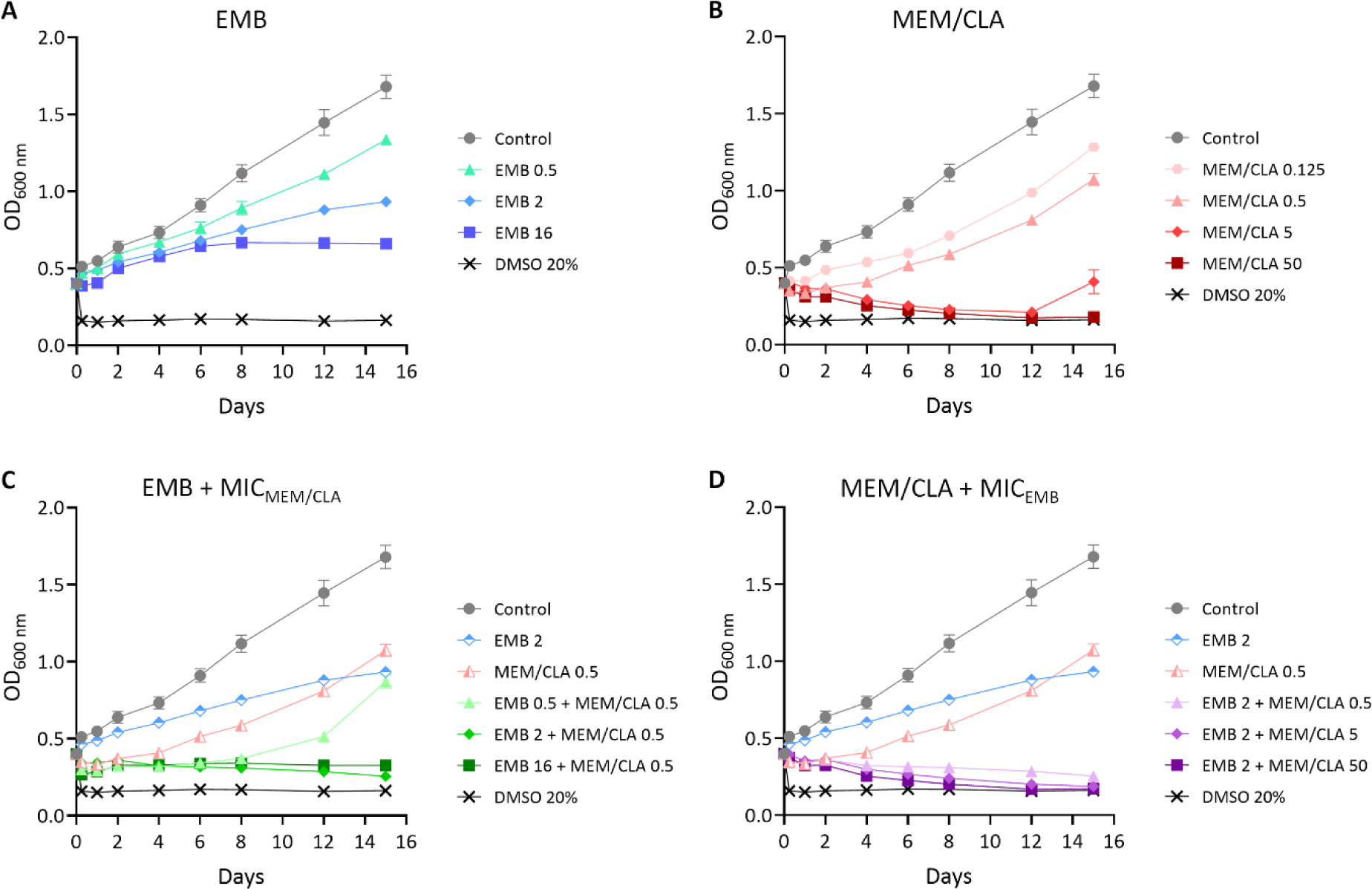
Growth curves of *Mycobacterium tuberculosis* (*Mtb*) H37Rv untreated or treated with different antibiotic conditions. (A) Treatment with increasing concentrations of ethambutol (EMB). (B) Treatment with increasing concentrations of meropenem/clavulanate (MEM/CLA). (C) Treatment with increasing concentrations of EMB and a fixed MEM/CLA concentration (MIC_MEM/CLA_ for *Mtb* H37Rv = 0.5 mg/L). (D) Treatment with increasing concentrations of MEM/CLA and a fixed EMB concentration (MIC_EMB_ for *Mtb* H37Rv = 2 mg/L). Optical density at 600 nm (OD_600_) was measured over 15 days for all cultures. Symbols show the mean value of at least three biological replicates and error bars represent standard error of the mean. MIC, minimum inhibitory concentration. The numbers after each antibiotic abbreviature indicate the respective concentration in mg/L. When present, clavulanate concentration was fixed at 2.5 mg/L.

Next, we observed the effects of combining MEM/CLA at the MIC with increasing concentrations of EMB (Figure 7C). In this case, all concentrations decreased the density, but after day 8, growth in the presence of the association with 0.5 mg/L of EMB started to increase again. This shift probably results from the inability of this concentration to achieve the full effect of the other concentrations, as observed in Figure 7A. Finally, when we fixed the concentration of EMB at the MIC and combined it with increasing MEM/CLA concentrations (Figure 7D), all the combinations resulted in similar density reductions. Importantly, the addition of EMB to 0.5 mg/L of MEM/CLA dramatically prolonged the effect of this concentration over time and produced the same effect as 50 mg/L MEM/CLA individually. Spots from the different treatments confirmed the outputs of the growth curves (Figure 8), with one exception noticed for 16 mg/L of EMB, which had a relatively high density, but no growth in the respective spot. This possibly results from the strong inhibition of the bacilli through a mechanism that does not involve lysis. Overall, the findings of this section indicate that EMB mostly potentiates the effects of meropenem over time without affecting the initial lytic capacity of the carbapenem.

**Figure 8.**
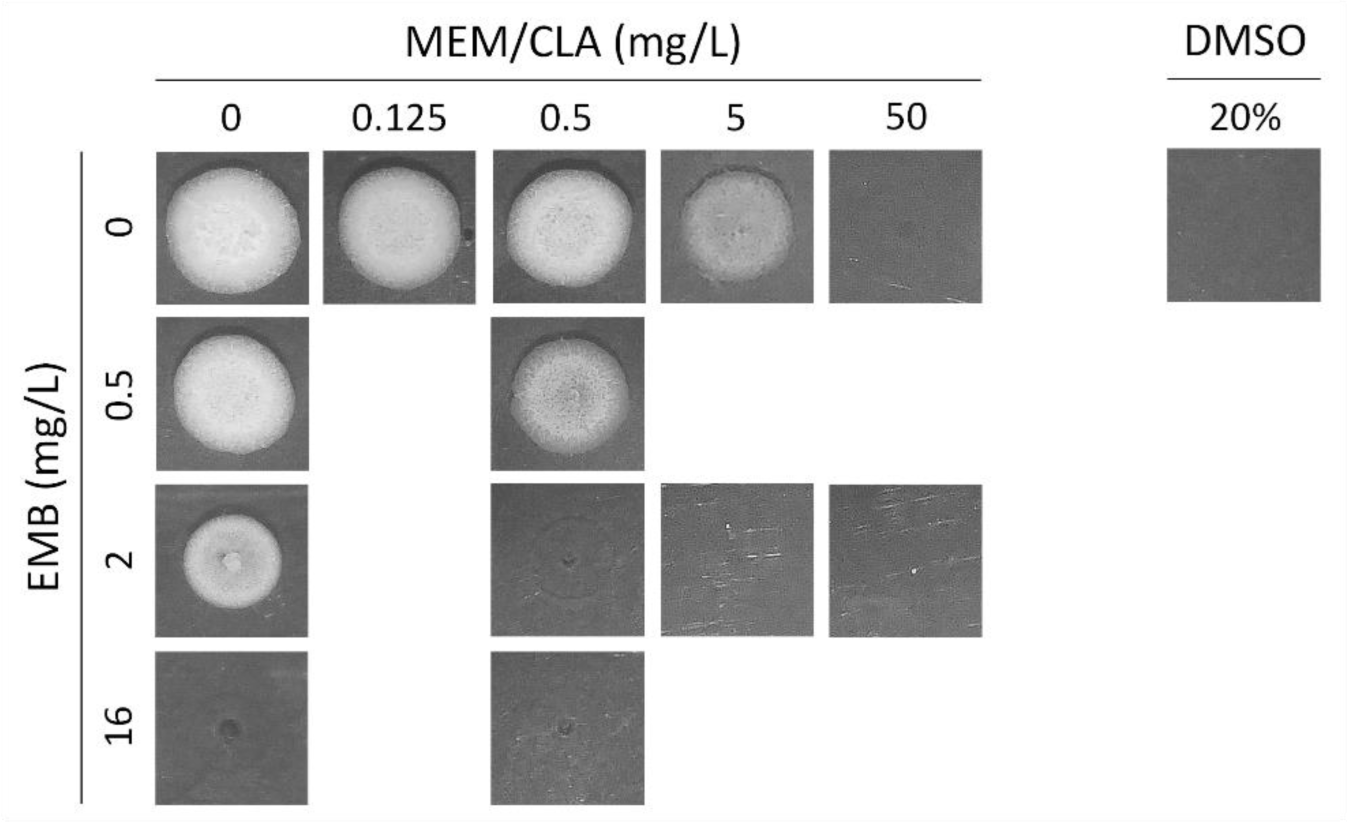
Spots of *Mycobacterium tuberculosis* H37Rv obtained from the growth assays with the antibiotics. After 15 days of incubation with the antibiotics (Figure 7), cultures were platted on antibiotic-free 7H10 agar plates and incubated until growth of the untreated control (7 days). The experiment was repeated in triplicate giving comparable results. CLA, clavulanate; EMB, ethambutol; MEM, meropenem.

## Discussion

Multiple studies in the last decades have rekindled the interest in applying beta-lactams against TB, specifically through repurposing of carbapenems (10,23,29,34–36). Since TB treatment requires complex schemes composed by multiple drugs for several months, determining potential interactions established between beta-lactams and other antibiotics is important to anticipate therapeutic outcomes. In this sense, interactions with INH and EMB, two first-line drugs that also target the cell wall, remain largely unknown, especially in clinical strains. In this work, we investigated these interactions in a set of *Mtb* strains, including the reference H37Rv and eight clinical strains with diverse resistance patterns and sublineage backgrounds. In addition, we explored if the most promising interaction was maintained intracellularly and if it had any impact in improved *Mtb* recognition by the host surveillance system.

EMB does not affect synthesis of MA but was shown to interfere with their transfer into the CW in *M. smegmatis* (37). Studies with *C. glutamicum* in which synthesis of AG was targeted through gene deletion or EMB treatment revealed that these decreased CW-bound MA and arabinose content (13,38). Given these observations, we expected that co-treatment with EMB and INH would result in a synergistic effect on the strains. However, this produced additive effects at most. Conversely, a synergistic minimum FICI was observed for the reference strain and for half of the clinical isolates when exposed to EMB combinations with beta-lactams. This is compatible with an earlier report that describes that subinhibitory concentrations of EMB reduced the MIC of AMX in *M. tuberculosis* isolates (39). It is also in agreement with synergies between EMB and penicillins described for *C. glutamicum* and *M. phlei* (13). In this study, Schubert et al. demonstrated that the bacteriostatic effect of EMB derives from a selective arrest of apical cell wall synthesis. On the other hand, cell division seems to be EMB-insensitive and nascent PG continues to be inserted into newly formed septa. Amoxicillin and meropenem form stable complexes with *M. tuberculosis* cell-division protein FtsI (also designated PBP3 or PbpB) and inhibition of this enzyme by beta*-*lactam antibiotics blocks *Mtb* replication (40,41). Thus, the addition of beta-lactams that interfere with cell division and PG synthesis/hydrolysis balance to bacteria with an EMB-compromised elongation machinery adds to this challenge and probably contributes to a more efficient killing. It has also been proposed that an approach that combines inhibitors of both PBPs and Ldts might be more effective at inhibiting PG synthesis and at suppressing resistance emergence (7,42). In our work, while growth reduction effects were commonly observed with amoxicillin plus meropenem, a synergistic FICI was never reached. These findings emphasize the importance of using different classes of antibiotics against *Mtb* and imply that combining beta-lactams with EMB might be the most effective cell-wall targeting strategy among the possible associations. Notably, all clinical strains for which an EMB and MEM/CLA synergy was reported were members of sublineage 4.3.4.2, which includes spoligotypes Latin American and Mediterranean (LAM) 1, LAM 4, and LAM11. In a previous work, we have shown that strains from this sublineage had significantly lower beta-lactam MICs and an earlier study also reported overrepresentation of AMX/CLA susceptibility among LAM4 strains (10,23). Our observation that EMB enhances the activity of meropenem against this sublineage again reinforces the hypothesis that LAM isolates are more susceptible to beta-lactams than other strains.

We then focused on the combination of MEM/CLA with EMB and evaluated its effects intracellularly. In conditions containing only the carbapenem, the pathogen eventually overcame its effects. Excluding the susceptible strains, the same applied to EMB. While the associations elicited a decline in intracellular growth that generally did not surpass the 2-log threshold, our results suggest that these treatments consistently ensured a faster and more sustained reduction of the intracellular bacilli, or at least, a static effect in the resistant strains treated with a subinhibitory concentration. This highlights that the level of resistance to EMB may limit the extent of the effects of this combination. Previous studies investigating the penetration of meropenem into pneumonic human lung tissue after intravenous administration of 1 g three times daily have determined a maximum interstitial concentration of 11.4 mg/L, which allows an effective killing of susceptible extracellular bacteria (43). Activity of beta-lactams against intraphagosomal organisms like *Mtb* is generally perceived as lower since drug accumulation in the cytosol is prevented and concentration within these vesicles is anticipated to be even lower (21,44,45). In an THP-1 infection model with the intraphagosomal bacteria *Listeria monocytogenes*, Lemaire et al. proposed that poor cellular accumulation may be offset by increased extracellular concentrations and prolonged exposure, leading to significant intracellular bacterial load reductions (46). This hypothesis resonates with the findings of our infection model and may explain why the most significant differences of the combinations compared to the individual antibiotics tended to emerge from combinations with the two lowest MEM concentrations. Regarding the intracellular activity of 50 mg/L meropenem, which roughly corresponds to the average maximum serum or plasma concentrations achieved after intravenous administration of 1 g (43,47), it seemed to already produce a considerable *Mtb* inhibition independently and adding EMB did not significantly improve on its effect.

In *C. glutamicum*, EMB mostly interferes with the acquisition of the polar and septal mycolate-layer, while sparing the side-wall content (48). This interference may contribute to increased polar PG exposure, and while BODIPY FL fluorescence following EMB exposure was only significantly higher in one strain, combination with MEM delivered fluorescence increases for all strains. As previously reported in both *Mtb* and *M. smegmatis*, BODIPY FL vancomycin binding occurred predominantly at the poles of the bacilli (28,49). Vancomycin recognizes the terminal d-alanyl-d-alanine moiety present in the PG precursors, which is also the target of DD-carboxypeptidases, enzymes that generate the substrate used by Ldts to form 3-3 cross-links (50). Besides its canonical action against PG transpeptidases, meropenem can additionally inhibit DD-carboxypeptidases and lead to pentapeptide stem accumulation (33). This increases the available binding sites for vancomycin and justifies the improved signal of its fluorescent analogue after incubation with MEM. Hence, we speculate that our observations with the combination may be the consequence of an EMB-enabled CW destabilization that enhances the access of the carbapenem to the periplasmic space where the PG transpeptidases locate and operate (8). Treatments of strain MDR III with a subinhibitory concentration of EMB produced the same BODIPY FL fluorescence intensities as incubations with EMB at the MIC. This suggests that resistance to EMB did not largely impact its ability to destabilize the mycobacterial CW during the period of the assay.

Despite the increase in CW destabilization and nascent PG exposure, this did not translate in a stronger macrophage inflammatory response. Treatments were added post-infection and it is possible that their impact in initial recognition by cell surface receptors was limited. As the infection progresses and some bacteria escape necrotic infected macrophages, the bacilli may encounter a more hostile environment that could enhance their recognition if both EMB and MEM are present. However, the lack of differences in IL-1β secretion among macrophages infected with pre-treated bacteria supports that the combination does not lead to improved extracellular pathogen recognition compared to MEM/CLA individually. A limitation of using BODIPY FL vancomycin is that it binds to stem peptide residues that may not overlap with the motifs recognized by some PRRs. Nonetheless, if co-treatments induced significant changes in the amount of PRR agonists, we expect this would have probably been captured by the highly sensitive ELISA technique. Therefore, in this study, IL-1β secretion levels seem to be strictly correlated with MEM/CLA concentration only. Treatment of *Mtb* with MEM/CLA results in rapid cell lysis accompanied by extravasation of cytoplasmic contents (33). Our results in bacteria growing in broth medium show that MEM/CLA reduces the density of bacterial suspensions, but this effect is weakened over time for concentrations below 50 mg/L if EMB is not simultaneously available. Significantly, concurrent presence of EMB and MEM/CLA at the MIC values prompted the same decrease in absorbance as a 100-fold higher carbapenem concentration applied individually. This expresses that lytic effects of the combination over extracellular bacteria are intense and sustained over a long period. However, the several tested antibiotic conditions do not seem to promote considerable differences between each other on the first day of incubation. It must be highlighted that in these assays, the bacterial suspensions are much denser than the initially applied in the infection assays and the amount of available antibiotics per bacteria was possibly lower. The dynamics within the macrophages may be very different, with the cells grating protection to lower MEM concentrations and partially metabolizing the antibiotics (46,51). Thus, it is reasonable to consider that suprainhibitory concentrations of MEM diffuse through the cells more efficiently and overcome some of these limitations, leading to a more extensive PG crosslinking inhibition and an increased intracellular pathogen lysis with release of PG fragments or mycobacterial nucleic acids (33).

Recognition of the released PAMPs by intracellular PRRs, including NOD2, NLRP3, and AIM2, triggers NFκB-dependent and inflammasome-mediated pro-inflammatory cytokine production (52–55). In accordance with an earlier study (56), we did not observe signs that therapeutically relevant MEM concentrations negatively affected cell viability over time, but the addition of the higher concentration of the carbapenem during infection with strain MDR III did induce a higher degree of macrophage necrosis. During *Mtb* infection, diverse cell death modes can be initiated, including inflammasome-mediated pyroptosis (57). During this particular type of necrotic cell death, highly inflammatory immune responses are promoted, namely IL-1β secretion via pore forming protein gasdermin D and release of damage-associated molecular patterns (DAMPs) (58–60). These facts together can feasibly explain the increase in necrotic cell proportion and the sharp rise in IL-1β secretion attained in the presence of higher MEM/CLA concentrations. Necrotic cells decreased when 16 mg/L of EMB were used, and this shift was associated with decreased IL-1β secretion during macrophage coincubation with 5 mg/L MEM/CLA. To a lesser extent, this decrease was also noticed for the antibiotic combinations in strain Susceptible I and might be attributed to the non-lytic inhibition of EMB (13). IL-1β is a critical cytokine during immune response to *Mtb* infection and the pathogen has been shown to inhibit the inflammasome–mediated IL-1β production through the ESX-1 secretion system (61). Moreover, in highly inflammatory host responses, as those involving the release of nitric oxide (NO), a controlled activation of the NLRP3 inflammasome was observed (62). NO-controlled NLRP3 activation was shown to be an evasion strategy to prevent exaggerated neutrophil recruitment to the granuloma, thereby sustaining Mtb survival in the host (63,64). Furthermore, isolates associated with severe TB induce lower IL-1β responses than those from mild cases, suggesting that this inflammatory cytokine can provide information on TB severity outcomes (65). Strain MDR III displayed a much lower IL-1β concentration when compared to the other two strains, which was completely abolished upon treatment with a MEM/CLA concentration above the MIC. Since low-IL-1β-inducing *Mtb* clinical isolates can evade the inflammasome (65), the observation that meropenem exposure may greatly restrict the pathogen’s ability to avoid this surveillance system is promising.

The present study confirmed that EMB establishes a synergy with beta-lactams against *Mtb* clinical isolates. Intracellularly, interaction of EMB with MEM/CLA potentially offers macrophages a temporal advantage to clear the pathogen. Considering beta-lactams are well known for a bactericidal activity heavily reliant on the time for which antibiotic levels exceed the MIC (66), combining them with drugs that allow a more intense or sustained effect may be pivotal to boost their activity (Figure 9). Since outer layers of the mycobacterial cell envelope are anchored to the peptidoglycan (1), we cannot exclude that beta-lactam-mediated inhibition of this polymer may also impact EMB activity over AG synthesis. We must equally consider the hypothesis that these antibiotics may influence host responses. Here, we have revealed that meropenem induces an exacerbated IL-1β secretion by *Mtb*-infected THP-1 macrophages. This finding should be further explored since host-directed therapies that modulate this cytokine may influence TB progression (67,68). Overall, our outputs contribute to clarifying the placement of beta-lactams in alternative regimens against *Mtb*, in alignment with WHO requirements for this class (9). Some constraints to clinical application of this class include the intravenous administration of carbapenems, which may be circumvented with the oral tebipenem (69,70). Another limitation is the low activity of beta-lactam/beta-lactam inhibitor combinations against nonreplicating *Mtb* (34), but novel cephalosporins that selectively act on this population have been identified (71,72). Finally, interactions between beta-lactams and anti-TB drugs other than ethambutol and isoniazid should be determined in greater detail. CW synthesis inhibitors cause an ATP burst associated with lethal effects in *Mycobacterium bovis* (73) and co-treatment of *M. abscessus* with the ATP synthase inhibitor bedaquiline eliminates the bactericidal activity of beta-lactams (74). Thus, clarifying possible antagonisms between beta-lactams and other anti-TB drugs in *Mtb* is especially urgent to avoid counterproductive therapeutic outcomes.

**Figure 9.**
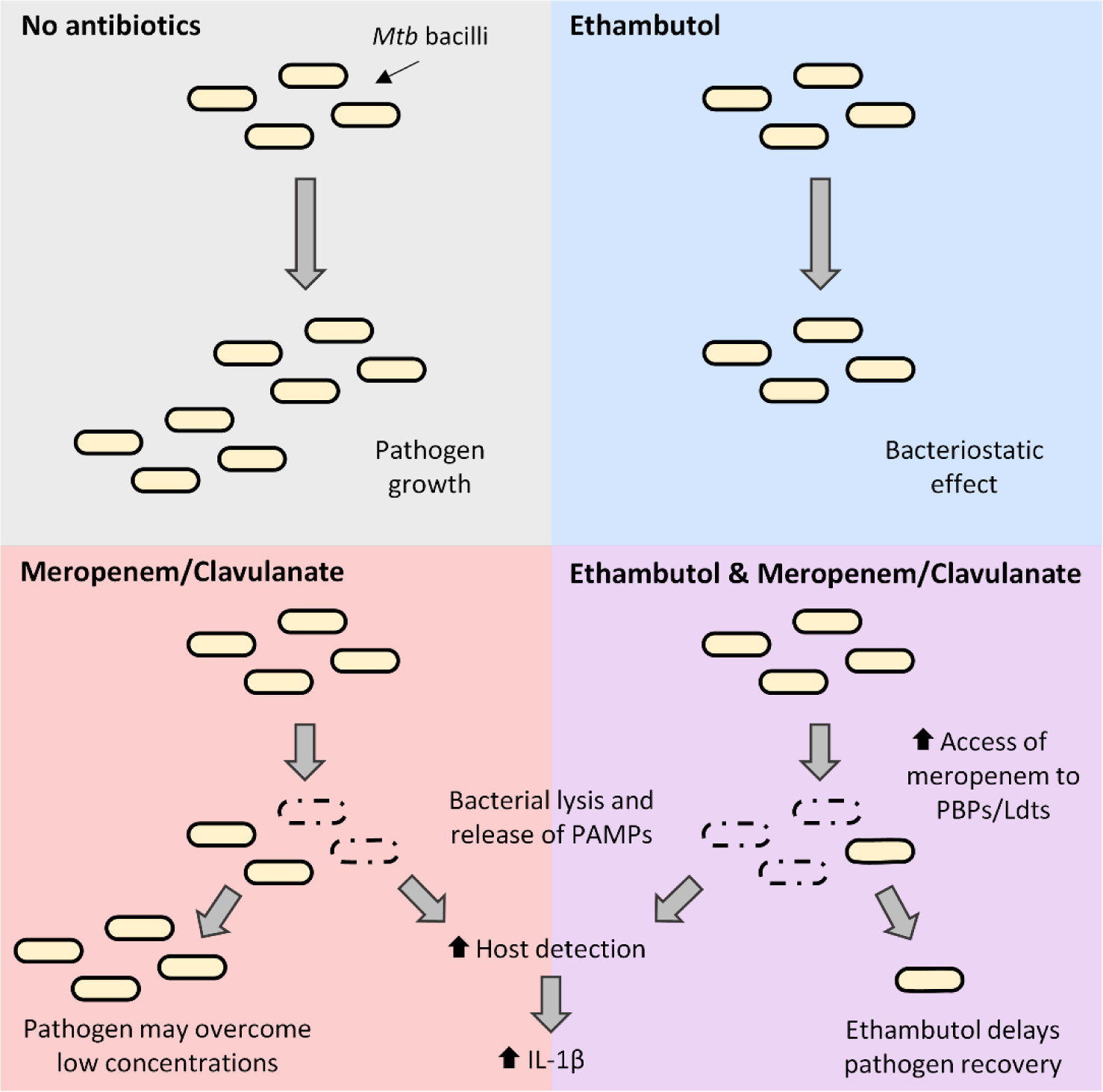
Schematic summary representation of the individual and combined effects of ethambutol and meropenem in this study. In the absence of antibiotics, *Mycobacterium tuberculosis* (*Mtb*) is able to grow normally. Ethambutol exerts a bacteriostatic effect, but some bacteria may still replicate slowly. Intracellularly, the agent allows faster elimination of ethambutol-susceptible strains. Meropenem/clavulanate promotes bacterial lysis and extravasation of cellular content, which can be sensed by host macrophages and induce IL-1β secretion. Surviving bacilli may overcome low concentrations of the carbapenem and result in the recovery of the pathogen. Combination with ethambutol increases the access of meropenem to peptidoglycan transpeptidases and inhibits the recovery of the pathogen after exposure to the carbapenem. This potentially enables macrophages to clear the infection more efficiently. Ldts, LD-transpeptidases; PBPs, penicillin-binding proteins.

## Author Contributions

FO, EA, and MJC conceived and designed the study. FO, DP, CS, BG, and FH performed the experiments. FO, DP, EA, and MJC analyzed the data. FO wrote the initial draft of the manuscript. EA and MJC revised and edited the manuscript. All authors reviewed and approved the final version of the manuscript.

## Acknowledgments

The authors would like to thank Catarina Godinho Santos and the Flow Cytometry Facility of iMed.ULisboa - Research Institute for Medicines for their services and assistance. The authors would also like to acknowledge Rita Macedo and the National Reference Laboratory for Mycobacteria - Instituto Nacional de Saúde Doutor Ricardo Jorge for providing the clinical strains.

## Supplementary Material

Supplementary Table S1 and Figures S1-S11 are provided in PDF format.

## References

1. Abrahams KA, Besra GS. Mycobacterial cell wall biosynthesis: a multifaceted antibiotic target. Parasitology. 2018;145(2):116–33.

2. Jankute M, Cox JAG, Harrison J, Besra GS. Assembly of the Mycobacterial Cell Wall. Annu Rev Microbiol. 2015;69:405–23.

3. Brennan PJ, Nikaido H. The envelope of mycobacteria. Annu Rev Biochem. 1995;64:29–63.

4. Abraham EP, Chain E, Fletcher CM, Gardner AD, Heatley NG, Jennings MA, et al. Further observations on penicillin. Lancet. 1941;238(6155):177–89.

5. Hugonnet JE, Tremblay LW, Boshoff HI, Barry CE 3rd, Blanchard JS. Meropenem-Clavulanate Is Effective Against Extensively Drug-Resistant Mycobacterium tuberculosis. Science. 2009;323(5918):1215–8.

6. Gupta R, Lavollay M, Mainardi JL, Arthur M, Bishai WR, Lamichhane G. The *Mycobacterium tuberculosis* protein Ldt_Mt2_ is a nonclassical transpeptidase required for virulence and resistance to amoxicillin. Nat Med. 2010;16(4):466–9.

7. Story-Roller E, Lamichhane G. Have we realized the full potential of β-lactams for treating drug-resistant TB? IUBMB Life. 2018;70(9):881–8.

8. Catalão MJ, Filipe SR, Pimentel M. Revisiting Anti-tuberculosis Therapeutic Strategies That Target the Peptidoglycan Structure and Synthesis. Front Microbiol. 2019;10:190.

9. World Health Organization. WHO consolidated guidelines on tuberculosis. Module 4: treatment - drug-resistant tuberculosis treatment, 2022 update. 2022.

10. Olivença F, Nunes A, Macedo R, Pires D, Silveiro C, Anes E, et al. Uncovering Beta-Lactam Susceptibility Patterns in Clinical Isolates of *Mycobacterium tuberculosis* through Whole-Genome Sequencing. Microbiol Spectr. 2022;10(4):e0067422.

11. Olivença F, Ferreira C, Nunes A, Silveiro C, Pimentel M, Gomes JP, et al. Identification of drivers of mycobacterial resistance to peptidoglycan synthesis inhibitors. Front Microbiol. 2022;13:985871.

12. Kaushik A, Makkar N, Pandey P, Parrish N, Singh U, Lamichhane G. Carbapenems and Rifampin Exhibit Synergy against *Mycobacterium tuberculosis* and *Mycobacterium abscessus*. Antimicrob Agents Chemother. 2015;59(10):6561–7.

13. Schubert K, Sieger B, Meyer F, Giacomelli G, Böhm K, Rieblinger A, et al. The Antituberculosis Drug Ethambutol Selectively Blocks Apical Growth in CMN Group Bacteria. mBio. 2017;8(1):e02213–16.

14. Silveiro C, Marques M, Olivença F, Pires D, Mortinho D, Nunes A, et al. CRISPRi-mediated characterization of novel anti-tuberculosis targets: Mycobacterial peptidoglycan modifications promote beta-lactam resistance and intracellular survival. Front Cell Infect Microbiol. 2023;13:1089911.

15. Chamaillard M, Hashimoto M, Horie Y, Masumoto J, Qiu S, Saab L, et al. An essential role for NOD1 in host recognition of bacterial peptidoglycan containing diaminopimelic acid. Nat Immunol. 2003;4(7):702–7.

16. Girardin SE, Boneca IG, Viala J, Chamaillard M, Labigne A, Thomas G, et al. NOD2 Is a General Sensor of Peptidoglycan through Muramyl Dipeptide (MDP) Detection. J Biol Chem. 2003;278(11):8869–72.

17. Coulombe F, Divangahi M, Veyrier F, de Léséleuc L, Gleason JL, Yang Y, et al. Increased NOD2-mediated recognition of *N*-glycolyl muramyl dipeptide. J Exp Med. 2009;206(8):1709–16.

18. Lavollay M, Arthur M, Fourgeaud M, Dubost L, Marie A, Veziris N, et al. The Peptidoglycan of Stationary-Phase *Mycobacterium tuberculosis* Predominantly Contains Cross-Links Generated by L,D-Transpeptidation. J Bacteriol. 2008;190(12):4360–6.

19. Kanneganti TD, Lamkanfi M, Núñez G. Intracellular NOD-like Receptors in Host Defense and Disease. Immunity. 2007;27(4):549–59.

20. Stamm CE, Collins AC, Shiloh MU. Sensing of *Mycobacterium tuberculosis* and consequences to both host and bacillus. Immunol Rev. 2015;264(1):204–19.

21. Renard C, Vanderhaeghe HJ, Claes PJ, Zenebergh A, Tulkens PM. Influence of conversion of penicillin G into a basic derivative on its accumulation and subcellular localization in cultured macrophages. Antimicrob Agents Chemother. 1987;31(3):410–6.

22. World Health Organization. Global Tuberculosis Report 2022. 2022.

23. Cohen KA, El-Hay T, Wyres KL, Weissbrod O, Munsamy V, Yanover C, et al. Paradoxical Hypersusceptibility of Drug-resistant *Mycobacterium tuberculosis* to β-lactam Antibiotics. EBioMedicine. 2016;9:170–9.

24. Omollo C, Singh V, Kigondu E, Wasuna A, Agarwal P, Moosa A, et al. Developing Synergistic Drug Combinations To Restore Antibiotic Sensitivity in Drug-Resistant *Mycobacterium tuberculosis*. Antimicrob Agents Chemother. 2021;65(5):e02554–20.

25. Noel DJ, Keevil CW, Wilks SA. Synergism versus Additivity: Defining the Interactions between Common Disinfectants. mBio. 2021;12(5):e0228121.

26. Jordão L, Bleck CKE, Mayorga L, Griffiths G, Anes E. On the killing of mycobacteria by macrophages. Cell Microbiol. 2008;10(2):529–48.

27. Pires D, Bernard EM, Pombo JP, Carmo N, Fialho C, Gutierrez MG, et al. *Mycobacterium tuberculosis* Modulates miR-106b-5p to Control Cathepsin S Expression Resulting in Higher Pathogen Survival and Poor T-Cell Activation. Front Immunol. 2017;8:1819.

28. Jeon AB, Obregón-Henao A, Ackart DF, Podell BK, Belardinelli JM, Jackson M, et al. 2-aminoimidazoles potentiate β-lactam antimicrobial activity against *Mycobacterium tuberculosis* by reducing β-lactamase secretion and increasing cell envelope permeability. PLoS One. 2017;12(7):e0180925.

29. Zhang D, Wang Y, Lu J, Pang Y. In Vitro Activity of β-Lactams in Combination with β-Lactamase Inhibitors against Multidrug-Resistant *Mycobacterium tuberculosis* Isolates. Antimicrob Agents Chemother. 2016;60(1):393–9.

30. Hugonnet JE, Blanchard JS. Irreversible Inhibition of the *Mycobacterium tuberculosis* β-Lactamase by Clavulanate. Biochemistry. 2007;46(43):11998–2004.

31. Ollinger J, Bailey MA, Moraski GC, Casey A, Florio S, Alling T, et al. A Dual Read-Out Assay to Evaluate the Potency of Compounds Active against *Mycobacterium tuberculosis*. PLoS One. 2013;8(4):e60531.

32. Ansel HC, Norred WP, Roth IL. Antimicrobial Activity of Dimethyl Sulfoxide Against *Escherichia coli*, *Pseudomonas aeruginosa*, and *Bacillus megaterium*. J Pharm Sci. 1969;58(7):836–9.

33. Kumar P, Arora K, Lloyd JR, Lee IY, Nair V, Fischer E, et al. Meropenem inhibits D,D-carboxypeptidase activity in *Mycobacterium tuberculosis*. Mol Microbiol. 2012;86(2):367–81.

34. Solapure S, Dinesh N, Shandil R, Ramachandran V, Sharma S, Bhattacharjee D, et al. In Vitro and In Vivo Efficacy of β-Lactams against Replicating and Slowly Growing/Nonreplicating *Mycobacterium tuberculosis*. Antimicrob Agents Chemother. 2013;57(6):2506–10.

35. Li F, Wan L, Xiao T, Liu H, Jiang Y, Zhao X, et al. In Vitro Activity of β-Lactams in Combination with β-Lactamase Inhibitors against *Mycobacterium tuberculosis* Clinical Isolates. Biomed Res Int. 2018;2018:3579832.

36. Gonzalo X, Satta G, Ortiz Canseco J, McHugh TD, Drobniewski F. Ertapenem and Faropenem against *Mycobacterium tuberculosis*: in vitro testing and comparison by macro and microdilution. BMC Microbiol. 2020;20(1):271.

37. Takayama K, Armstrong EL, Kunugi KA, Kilburn JO. Inhibition by Ethambutol of Mycolic Acid Transfer into the Cell Wall of *Mycobacterium smegmatis*. Antimicrob Agents Chemother. 1979;16(2):240–2.

38. Seidel M, Alderwick LJ, Birch HL, Sahm H, Eggeling L, Besra GS. Identification of a Novel Arabinofuranosyltransferase AftB Involved in a Terminal Step of Cell Wall Arabinan Biosynthesis in Corynebacterianeae, such as *Corynebacterium glutamicum* and *Mycobacterium tuberculosis*. J Biol Chem. 2007;282(20):14729–40.

39. Abate G, Miorner H. Susceptibility of multidrug-resistant strains of *Mycobacterium tuberculosis* to amoxycillin in combination with clavulanic acid and ethambutol. J Antimicrob Chemother. 1998;42(6):735–40.

40. Lu Z, Wang H, Zhang A, Liu X, Zhou W, Yang C, et al. Structures of *Mycobacterium tuberculosis* Penicillin-Binding Protein 3 in Complex with Five β-Lactam Antibiotics Reveal Mechanism of Inactivation. Mol Pharmacol. 2020;97(4):287–94.

41. Slayden RA, Belisle JT. Morphological features and signature gene response elicited by inactivation of FtsI in *Mycobacterium tuberculosis*. J Antimicrob Chemother. 2009;63(3):451–7.

42. Gonzalo X, Drobniewski F. Is there a place for β-lactams in the treatment of multidrug-resistant/extensively drug-resistant tuberculosis? Synergy between meropenem and amoxicillin/clavulanate. J Antimicrob Chemother. 2013;68(2):366–9.

43. Tomaselli F, Maier A, Matzi V, Smolle-Jüttner FM, Dittrich P. Penetration of Meropenem into Pneumonic Human Lung Tissue as Measured by In Vivo Microdialysis. Antimicrob Agents Chemother. 2004;48(6):2228–32.

44. Tulkens PM. Intracellular distribution and activity of antibiotics. Eur J Clin Microbiol Infect Dis. 1991;10(2):100–6.

45. Carryn S, Chanteux H, Seral C, Mingeot-Leclercq MP, Van Bambeke F, Tulkens PM. Intracellular pharmacodynamics of antibiotics. Infect Dis Clin North Am. 2003;17(3):615–34.

46. Lemaire S, Van Bambeke F, Mingeot-Leclercq MP, Tulkens PM. Activity of three β-lactams (ertapenem, meropenem and ampicillin) against intraphagocytic *Listeria monocytogenes* and *Staphylococcus aureus*. J Antimicrob Chemother. 2005;55(6):897–904.

47. Abulfathi AA, de Jager V, van Brakel E, Reuter H, Gupte N, Vanker N, et al. The Population Pharmacokinetics of Meropenem in Adult Patients With Rifampicin-Sensitive Pulmonary Tuberculosis. Front Pharmacol. 2021;12:637618.

48. Kumagai Y, Hirasawa T, Hayakawa K, Nagai K, Wachi M. Fluorescent Phospholipid Analogs as Microscopic Probes for Detection of the Mycolic Acid-Containing Layer in *Corynebacterium glutamicum*: Detecting Alterations in the Mycolic Acid-Containing Layer Following Ethambutol Treatment. Biosci Biotechnol Biochem. 2005;69(11):2051–6.

49. Joyce G, Williams KJ, Robb M, Noens E, Tizzano B, Shahrezaei V, et al. Cell Division Site Placement and Asymmetric Growth in Mycobacteria. PLoS One. 2012;7(9):e44582.

50. Pandey SD, Pal S, Kumar N G, Bansal A, Mallick S, Ghosh AS. Two DD-Carboxypeptidases from *Mycobacterium smegmatis* Affect Cell Surface Properties through Regulation of Peptidoglycan Cross-Linking and Glycopeptidolipids. J Bacteriol. 2018;200(14):e00760–17.

51. Diene SM, Pinault L, Keshri V, Armstrong N, Khelaifia S, Chabrière E, et al. Human metallo-β-lactamase enzymes degrade penicillin. Sci Rep. 2019;9(1):12173.

52. Mishra BB, Moura-Alves P, Sonawane A, Hacohen N, Griffiths G, Moita LF, et al. *Mycobacterium tuberculosis* protein ESAT-6 is a potent activator of the NLRP3/ASC inflammasome. Cell Microbiol. 2010;12(8):1046–63.

53. Brooks MN, Rajaram MVS, Azad AK, Amer AO, Valdivia-Arenas MA, Park JH, et al. NOD2 controls the nature of the inflammatory response and subsequent fate of *Mycobacterium tuberculosis* and *M. bovis* BCG in human macrophages. Cell Microbiol. 2011;13(3):402–18.

54. Saiga H, Kitada S, Shimada Y, Kamiyama N, Okuyama M, Makino M, et al. Critical role of AIM2 in *Mycobacterium tuberculosis* infection. Int Immunol. 2012;24(10):637–44.

55. Anes E, Pires D, Mandal M, Azevedo-Pereira JM. ESAT-6 a Major Virulence Factor of *Mycobacterium tuberculosis*. Biomolecules. 2023;13(6):968.

56. Matera G, Berlinghieri MC, Focà A. Meropenem: effects on human leukocyte functions and interleukin release. Int J Antimicrob Agents. 1995;5(2):129–33.

57. Chai Q, Wang L, Liu CH, Ge B. New insights into the evasion of host innate immunity by *Mycobacterium tuberculosis*. Cell Mol Immunol. 2020;17(9):901–13.

58. Shi J, Zhao Y, Wang K, Shi X, Wang Y, Huang H, et al. Cleavage of GSDMD by inflammatory caspases determines pyroptotic cell death. Nature. 2015;526(7575):660–5.

59. Evavold CL, Ruan J, Tan Y, Xia S, Wu H, Kagan JC. The Pore-Forming Protein Gasdermin D Regulates Interleukin-1 Secretion from Living Macrophages. Immunity. 2018;48(1):35–44.e6.

60. Beckwith KS, Beckwith MS, Ullmann S, Sætra RS, Kim H, Marstad A, et al. Plasma membrane damage causes NLRP3 activation and pyroptosis during *Mycobacterium tuberculosis* infection. Nat Commun. 2020;11(1):2270.

61. Shah S, Bohsali A, Ahlbrand SE, Srinivasan L, Rathinam VAK, Vogel SN, et al. Cutting Edge: *Mycobacterium tuberculosis* but Not Nonvirulent Mycobacteria Inhibits IFN-β and AIM2 Inflammasome–Dependent IL-1β Production via Its ESX-1 Secretion System. J Immunol. 2013;191(7):3514–8.

62. Mishra BB, Rathinam VAK, Martens GW, Martinot AJ, Kornfeld H, Fitzgerald KA, et al. Nitric oxide controls the immunopathology of tuberculosis by inhibiting NLRP3 inflammasome– dependent processing of IL-1β. Nat Immunol. 2013;14(1):52–60.

63. Mishra BB, Lovewell RR, Olive AJ, Zhang G, Wang W, Eugenin E, et al. Nitric oxide prevents a pathogen-permissive granulocytic inflammation during tuberculosis. Nat Microbiol. 2017;2:17072.

64. Dallenga T, Schaible UE. Neutrophils in tuberculosis – first line of defence or booster of disease and targets for host directed therapy? Pathog Dis. 2016;4(3):ftw012.

65. Sousa J, Cá B, Maceiras AR, Simões-Costa L, Fonseca KL, Fernandes AI, et al. *Mycobacterium tuberculosis* associated with severe tuberculosis evades cytosolic surveillance systems and modulates IL-1β production. Nat Commun. 2020;11(1):1949.

66. Turnidge JD. The Pharmacodynamics of β-Lactams. Clin Infect Dis. 1998;27(1):10–22.

67. Silvério D, Gonçalves R, Appelberg R, Saraiva M. Advances on the Role and Applications of Interleukin-1 in Tuberculosis. mBio. 2021;12(6):e0313421.

68. Moorlag SJCFM, Khan N, Novakovic B, Kaufmann E, Jansen T, van Crevel R, et al. β-Glucan Induces Protective Trained Immunity against *Mycobacterium tuberculosis* Infection: A Key Role for IL-1. Cell Rep. 2020;31(7):107634.

69. Horita Y, Maeda S, Kazumi Y, Doi N. In Vitro Susceptibility of *Mycobacterium tuberculosis* Isolates to an Oral Carbapenem Alone or in Combination with β-Lactamase Inhibitors. Antimicrob Agents Chemother. 2014;58(11):7010–4.

70. Gonzalo X, Drobniewski F. Are the Newer Carbapenems of Any Value against Tuberculosis. Antibiotics. 2022;11(8):1070.

71. Gold B, Smith R, Nguyen Q, Roberts J, Ling Y, Lopez Quezada L, et al. Novel Cephalosporins Selectively Active on Nonreplicating *Mycobacterium tuberculosis*. J Med Chem. 2016;59(13):6027–44.

72. Lopez Quezada L, Smith R, Lupoli TJ, Edoo Z, Li X, Gold B, et al. Activity-Based Protein Profiling Reveals That Cephalosporins Selectively Active on Non-replicating *Mycobacterium tuberculosis* Bind Multiple Protein Families and Spare Peptidoglycan Transpeptidases. Front Microbiol. 2020;11:1248.

73. Shetty A, Dick T. Mycobacterial cell wall synthesis inhibitors cause lethal ATP burst. Front Microbiol. 2018;9:1898.

74. Lindman M, Dick T. Bedaquiline Eliminates Bactericidal Activity of β-Lactams against *Mycobacterium abscessus*. Antimicrob Agents Chemother. 2019;63(8):e00827–19.

