## Supplementary Material for "Ethambutol and Meropenem/Clavulanate Synergy Promotes Enhanced Extracellular and Intracellular Killing of *Mycobacterium tuberculosis*"

Table S1. Information of the drug susceptibility testing and *in silico* TB-profiler predictions of lineages and spoligotypes of the *Mycobacterium tuberculosis* strains used in this study. The filled circle symbol (●) indicates resistance to the respective antibiotic. The dash symbol (-) indicates that the parameter was not assessed or that it does not apply. AMK, amikacin; AMX, amoxicillin; CLA, clavulanate; EMB, ethambutol; ETI, ethionamide; INH, isoniazid; KAN, kanamycin; LVX, levofloxacin; MEM, meropenem; MIC, minimum inhibitory concentration; MXF, moxifloxacin; PZA, pyrazinamide; RIF, rifampicin; STR, streptomycin; VAN, vancomycin.

| Strain | Strain ID | Drug-resistance profile | Antimicrobial susceptibility testing | | | | | | | | | | | | | | | TB Profiler *in silico* prediction | | Reference |
|---|---|---|---|---|---|---|---|---|---|---|---|---|---|---|---|---|---|---|---|---|
| | | | Antimycobacterial resistance (standardized guidelines) | | | | | | | | | | Broth microdilution assay MIC (mg/L) | | | | | | | |
| | | | INH | RIF | EMB | PZA | STR | LVX | MXF | AMK | KAN | ETI | INH | EMB | AMX/CLA | MEM/CLA | VAN | Sublineage | Spoligotype | |
| H37Rv WT | PT_Mtb001 | Susceptible | - | - | - | - | - | - | - | - | - | - | 0.031 | 2 | 2 | 0.5 | 16 | - | - | 11 |
| Susceptible I | PT_TB0245 | Susceptible | | | | | | | | | | | 0.031 | 1 | 1 | 0.5 | 16 | 4.1.2.1 | T1;H1 | 10 |
| Susceptible II | PT_TB0319 | Susceptible | | | | | | | | | | | 0.031 | 2 | 8 | 2 | 64 | 2.2.1.1 | Beijing-RD150 | 10 |
| Susceptible III | PT_TB0275 | Susceptible | | | | | | | | | | | 0.031 | 2 | 0.5 | 0.25 | 8 | 4.3.4.2 | LAM1;LAM4;LAM11 | 10 |
| Susceptible IV | PT_TB0283 | Susceptible | | | | | | | | | | | 0.063 | 4 | 0.5 | 0.5 | 8 | 4.3.4.2 | LAM1;LAM4;LAM11 | 10 |
| MDR I | PT_TB0022 | MDR | ● | ● | ● | | ● | | | | | ● | 4 | 8 | 0.25 | 0.25 | 32 | 4.3.4.2 | LAM1;LAM4;LAM11 | 10 |
| MDR II | PT_TB0055 | MDR | ● | ● | ● | ● | ● | | | | ● | ● | 32 | 16 | 1 | 0.5 | 2 | 4.2.1 | H3;H4 | 10 |
| MDR III | PT_TB0023 | MDR | ● | ● | ● | ● | ● | | | ● | ● | | 16 | 16 | 8 | 2 | 8 | 4.2.1 | H3;H4 | 10 |
| Pre-XDR | PT_TB0059 | Pre-XDR | ● | ● | ● | ● | ● | ● | ● | ● | | ● | 4 | 64 | 0.5 | 0.5 | 32 | 4.3.4.2 | LAM1;LAM4;LAM11 | 10 |

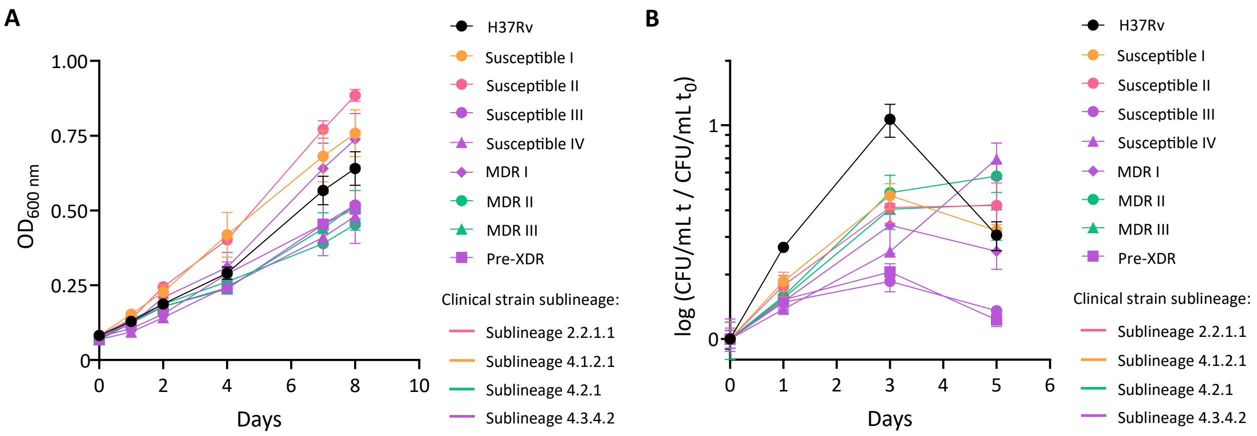


Figure S1. Growth profiles of *Mtb* H37Rv and of the eight clinical isolates used in this study. (A) Extracellular growth curves obtained by measuring the optical density at 600 nm ($OD_{600}$) of the bacterial suspensions over eight days. Symbols show the mean value of two replicates and error bars represent standard error of the mean. (B) Intracellular growth in a THP-1 infection model over five days of infection. The log of the ratio between the CFU/mL in each timepoint over the CFU/mL at day 0 is plotted. Symbols represent the average of at least 3 replicates per timepoint and error bars show the standard error of the mean. The color of the symbols denotes the respective sublineage of the clinical strains.

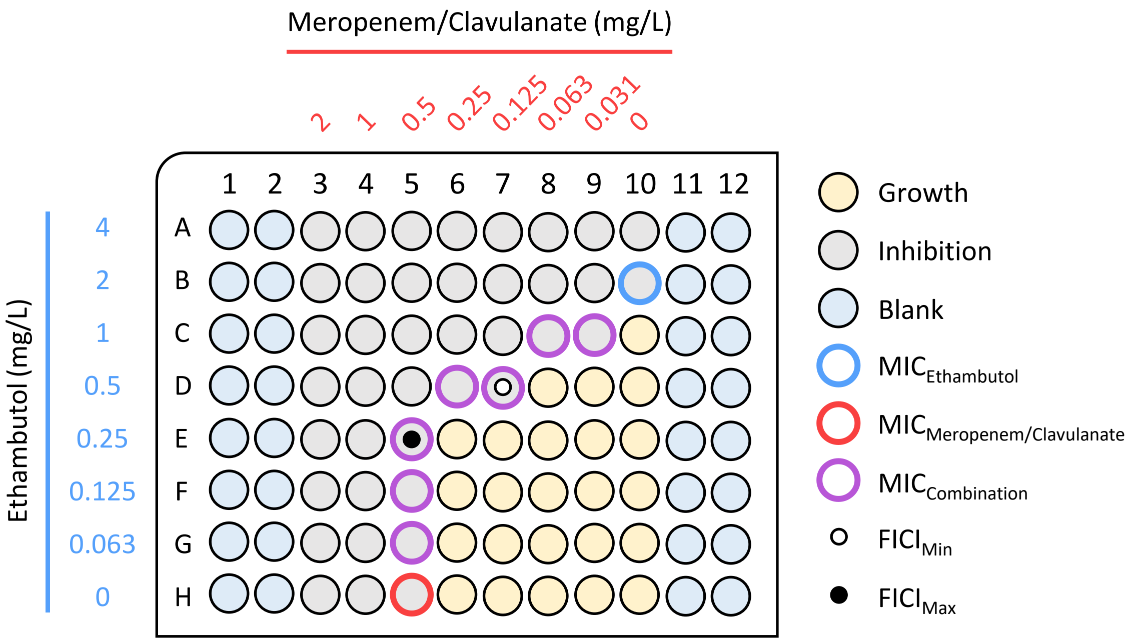


$$FICI_{Min} = \frac{0.5}{2} + \frac{0.125}{0.5} = 0.5$$

$$FICI_{Max} = \frac{0.25}{2} + \frac{0.5}{0.5} = 1.125$$

Figure S2. Schematic representation of the checkerboard assays employed in this study and of the calculation of the fractional inhibitory concentration (FIC) index (FICI) for the combination wells. The selected example depicts the results of one of the replicates of the combination between ethambutol and meropenem/clavulanate in *Mtb* H37Rv. $FICI_{Min}$, lowest FICI; $FICI_{Max}$, highest FICI; MIC, minimum inhibitory concentration. When present, clavulanate concentration was fixed at 2.5 mg/L.

| | | EMB & INH | EMB & AMX/CLA | EMB & MEM/CLA | INH & AMX/CLA | INH & MEM/CLA | AMX & MEM/CLA |
|---|---|---|---|---|---|---|---|
| H37Rv | $FICI_{Min}$ | 0.781 | 0.344 | 0.500 | 0.813 | 0.797 | 1.000 |
| | $FICI_{Med}$ | 1.063 | 0.578 | 0.750 | 1.438 | 1.250 | 1.125 |
| | $FICI_{Max}$ | 1.375 | 1.063 | 1.125 | 2.250 | 1.875 | 1.250 |
| Susceptible I | $FICI_{Min}$ | 1.031 | 0.594 | 0.625 | 1.094 | 1.000 | 1.063 |
| | $FICI_{Med}$ | 1.156 | 0.891 | 0.844 | 1.500 | 1.250 | 1.344 |
| | $FICI_{Max}$ | 1.375 | 1.188 | 1.141 | 2.250 | 2.125 | 2.188 |
| Susceptible II | $FICI_{Min}$ | 1.063 | 0.688 | 0.625 | 1.281 | 1.063 | 0.875 |
| | $FICI_{Med}$ | 1.625 | 1.063 | 1.000 | 2.063 | 1.375 | 1.125 |
| | $FICI_{Max}$ | 2.375 | 1.250 | 1.250 | 3.375 | 2.188 | 1.813 |
| Susceptible III | $FICI_{Min}$ | 0.563 | 0.375 | 0.375 | 0.531 | 0.547 | 0.563 |
| | $FICI_{Med}$ | 0.883 | 0.469 | 0.602 | 0.852 | 1.047 | 1.219 |
| | $FICI_{Max}$ | 1.156 | 0.797 | 1.078 | 2.688 | 2.188 | 2.000 |
| Susceptible IV | $FICI_{Min}$ | 0.875 | 0.438 | 0.375 | 0.844 | 0.578 | 0.875 |
| | $FICI_{Med}$ | 1.313 | 0.563 | 0.492 | 1.094 | 0.859 | 1.891 |
| | $FICI_{Max}$ | 3.125 | 0.625 | 0.594 | 2.000 | 1.000 | 2.500 |
| MDR I | $FICI_{Min}$ | 1.000 | 0.563 | 0.438 | 0.563 | 0.563 | 0.750 |
| | $FICI_{Med}$ | 1.156 | 0.750 | 0.563 | 0.750 | 1.000 | 0.906 |
| | $FICI_{Max}$ | 1.688 | 1.125 | 0.625 | 1.125 | 1.313 | 1.188 |
| MDR II | $FICI_{Min}$ | 0.781 | 0.500 | 0.563 | 0.766 | 0.781 | 0.625 |
| | $FICI_{Med}$ | 1.000 | 0.703 | 0.813 | 0.922 | 1.047 | 1.094 |
| | $FICI_{Max}$ | 1.625 | 1.625 | 1.188 | 1.688 | 1.688 | 1.625 |
| MDR III | $FICI_{Min}$ | 1.000 | 0.563 | 0.625 | 1.000 | 1.000 | 1.063 |
| | $FICI_{Med}$ | 1.250 | 0.875 | 1.000 | 1.188 | 1.125 | 1.250 |
| | $FICI_{Max}$ | 2.125 | 1.250 | 1.250 | 2.250 | 1.250 | 2.125 |
| Pre-XDR | $FICI_{Min}$ | 0.656 | 0.219 | 0.281 | 0.594 | 0.531 | 0.531 |
| | $FICI_{Med}$ | 0.906 | 0.313 | 0.406 | 1.250 | 0.672 | 0.625 |
| | $FICI_{Max}$ | 1.188 | 0.547 | 0.797 | 2.750 | 1.125 | 0.875 |

Type of Interaction: Synergy (FIC Index ≤ 0.5) Additive (0.5 < FIC Index ≤ 1) Indifferent (1 < FIC Index ≤ 4)

Figure S3. Heatmap of the detailed outputs of the checkerboard assays with six antibiotic combinations in *Mtb* H37Rv and eight clinical isolates. The results show the values of lowest fractional inhibitory concentration (FIC) index ($FICI_{Min}$), the median FICI ($FICI_{Med}$), and the highest FICI ($FICI_{Max}$) calculated as the mean of two independent replicates for each combination and in each strain. The color of the cells depicts the type of interaction based on the FICI value. AMX, amoxicillin; CLA, clavulanate; EMB, ethambutol; INH, isoniazid; MEM, meropenem.

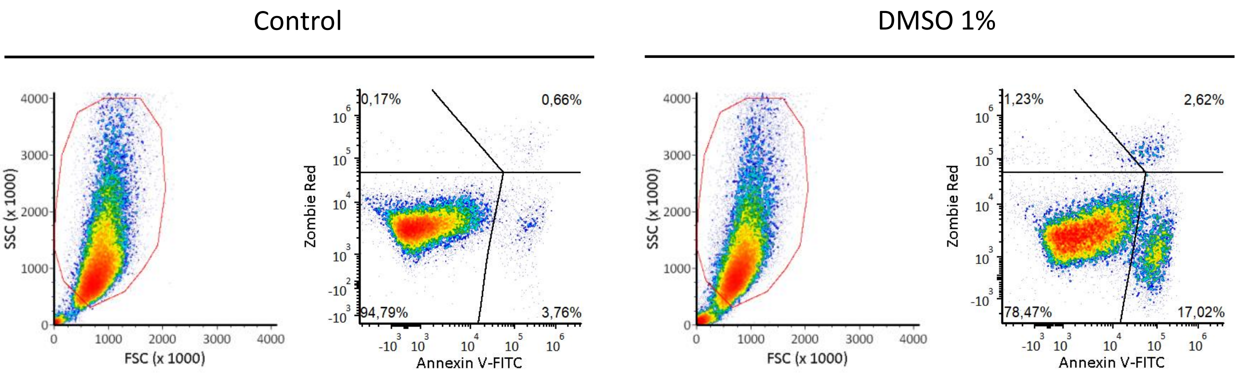


Figure S4. Representative replicates of the flow cytometry analysis of uninfected THP-1 macrophages present in Figure 4B. For each condition, the left panel shows the macrophage population gated for analysis and the right panel depicts the respective percentage of cells stained for annexin V and/or for Zombie Red on day 3 of treatment.

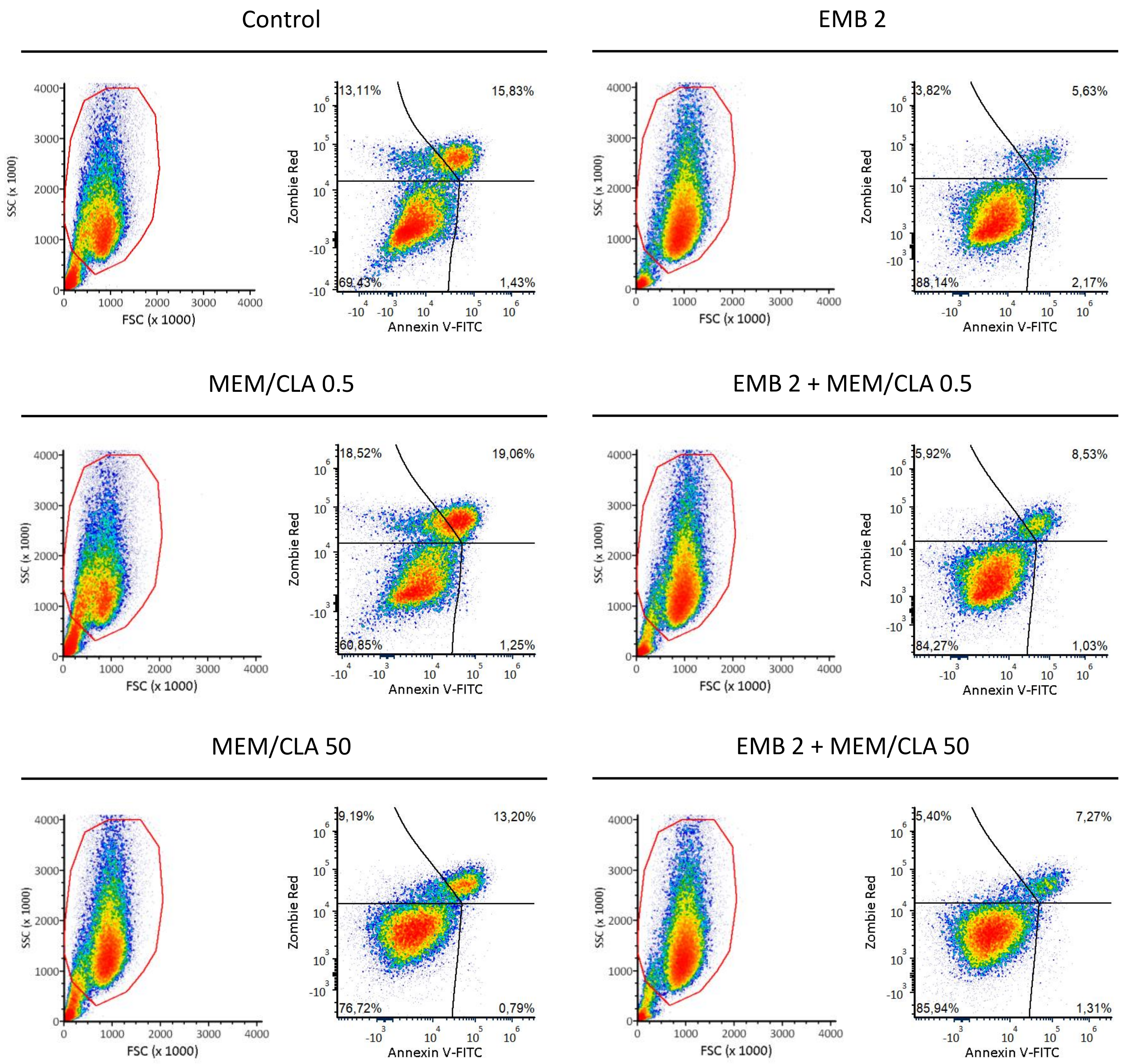


Figure S5. Representative replicates of the flow cytometry analysis of THP-1 macrophages infected with *Mycobacterium tuberculosis* H37Rv present in Figure 4B. For each condition, the left panel shows the macrophage population gated for analysis and the right panel depicts the respective percentage of cells stained for annexin V and/or for Zombie Red on day 3 post-infection. CLA, clavulanate; EMB, ethambutol; MEM, meropenem. The numbers after each antibiotic abbreviature indicate the respective concentration in mg/L. When present, clavulanate concentration was fixed at 2.5 mg/L.

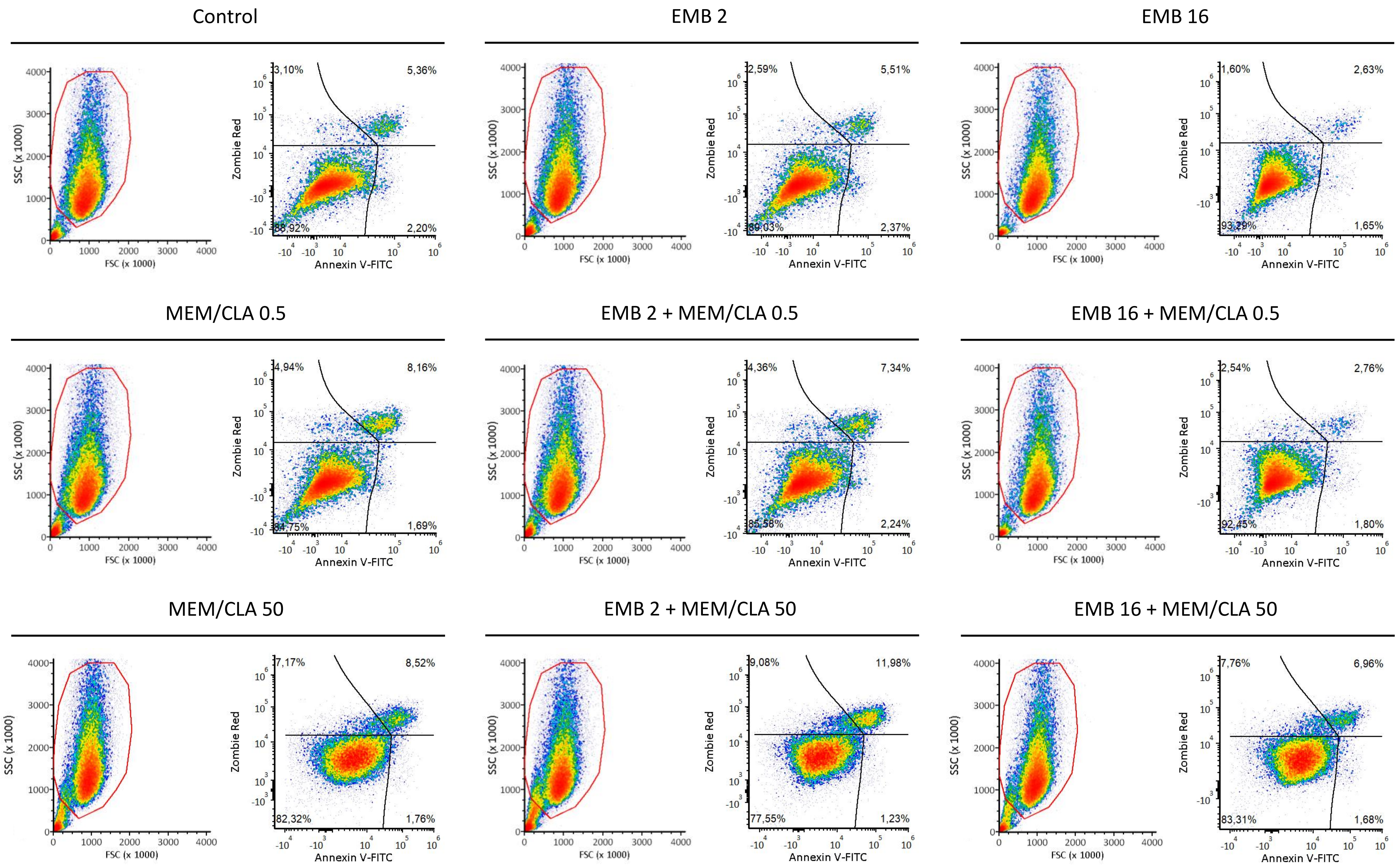


Figure S6. Representative replicates of the flow cytometry analysis of THP-1 macrophages infected with strain MDR III present in Figure 4B. For each condition, the left panel shows the macrophage population gated for analysis and the right panel depicts the respective percentage of cells stained for annexin V and/or for Zombie Red on day 3 post-infection. CLA, clavulanate; EMB, ethambutol; MEM, meropenem. The numbers after each antibiotic abbreviature indicate the respective concentration in mg/L. When present, clavulanate concentration was fixed at 2.5 mg/L.

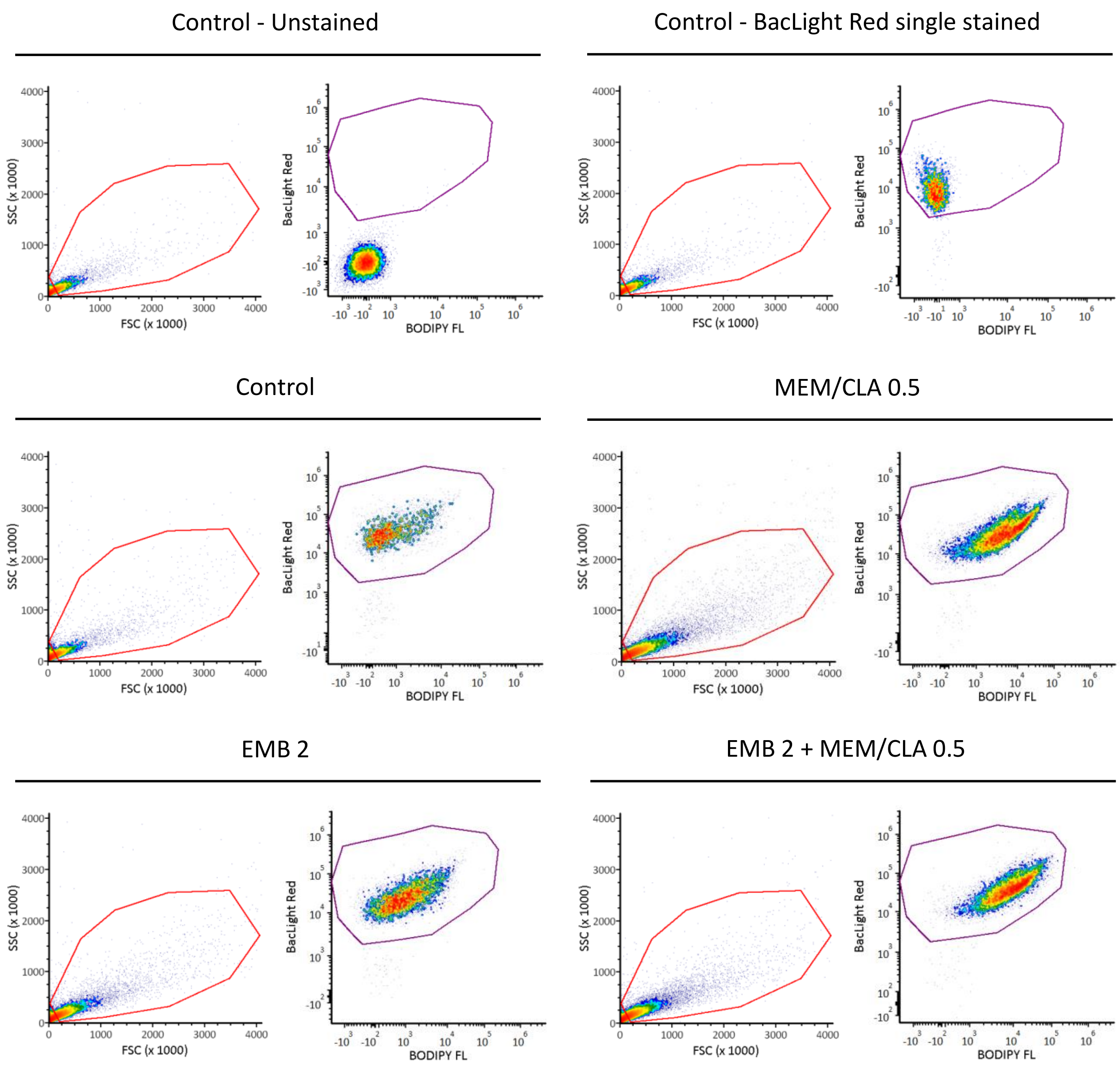


Figure S7. Representative replicates of the flow cytometry analysis of *Mycobacterium tuberculosis* strain H37Rv present in Figures 5A-B. For each condition, the left panel shows the bacterial population gated for analysis (red gate) and the right panel shows the distribution of bacteria stained for BacLight Red and/or for BODIPY FL after 6 h of treatment. BODIPY FL fluorescence intensity was quantified from bacteria stained with BacLight Red (purple gate). CLA, clavulanate; EMB, ethambutol; MEM, meropenem. The numbers after each antibiotic abbreviature indicate the respective concentration in mg/L. When present, clavulanate concentration was fixed at 2.5 mg/L.

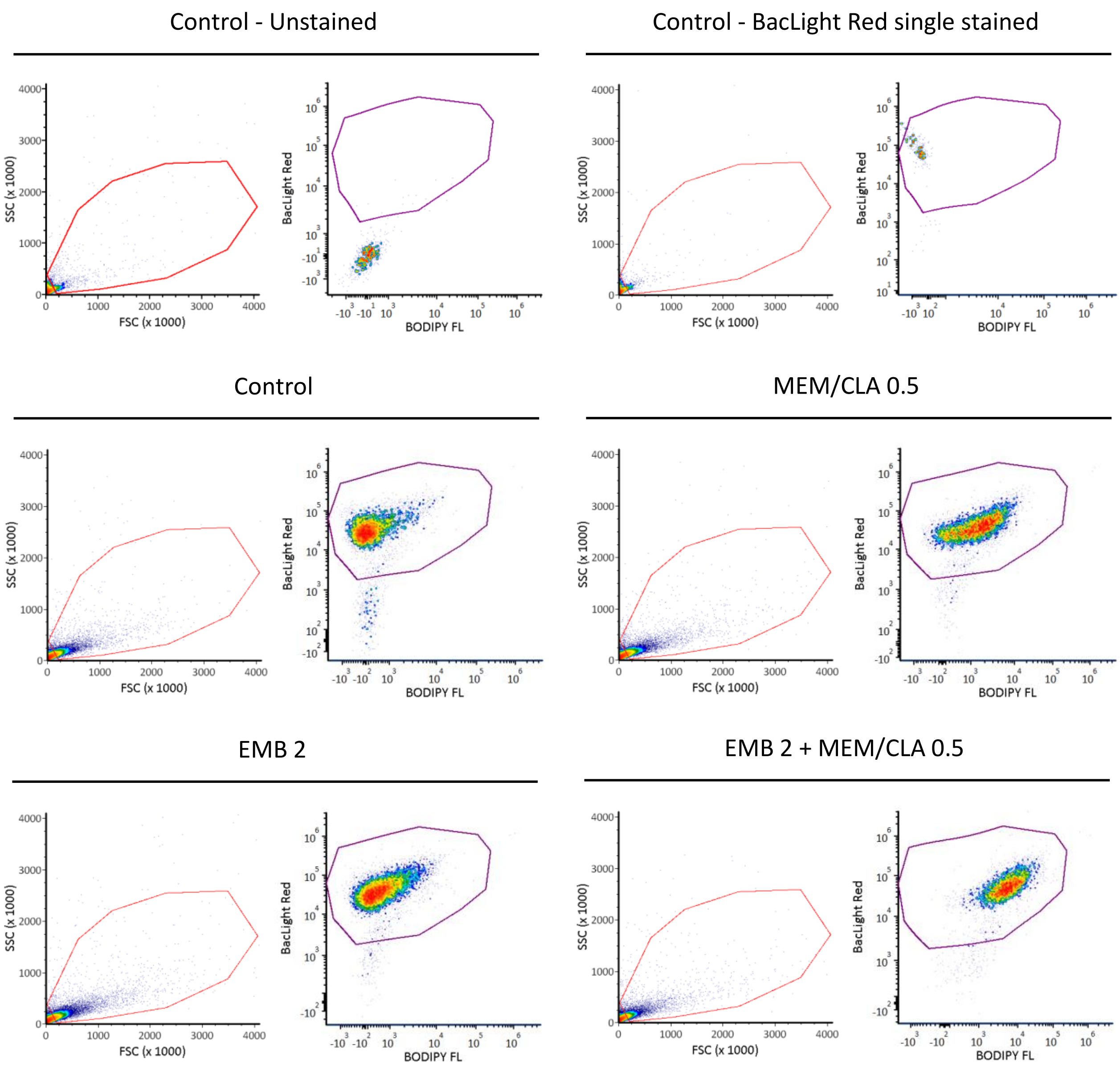


Figure S8. Representative replicates of the flow cytometry analysis of *Mycobacterium tuberculosis* strain Susceptible III present in Figures 5A-B. For each condition, the left panel shows the bacterial population gated for analysis (red gate) and the right panel shows the distribution of bacteria stained for BacLight Red and/or for BODIPY FL after 6 h of treatment. BODIPY FL fluorescence intensity was quantified from bacteria stained with BacLight Red (purple gate). CLA, clavulanate; EMB, ethambutol; MEM, meropenem. The numbers after each antibiotic abbreviature indicate the respective concentration in mg/L. When present, clavulanate concentration was fixed at 2.5 mg/L.

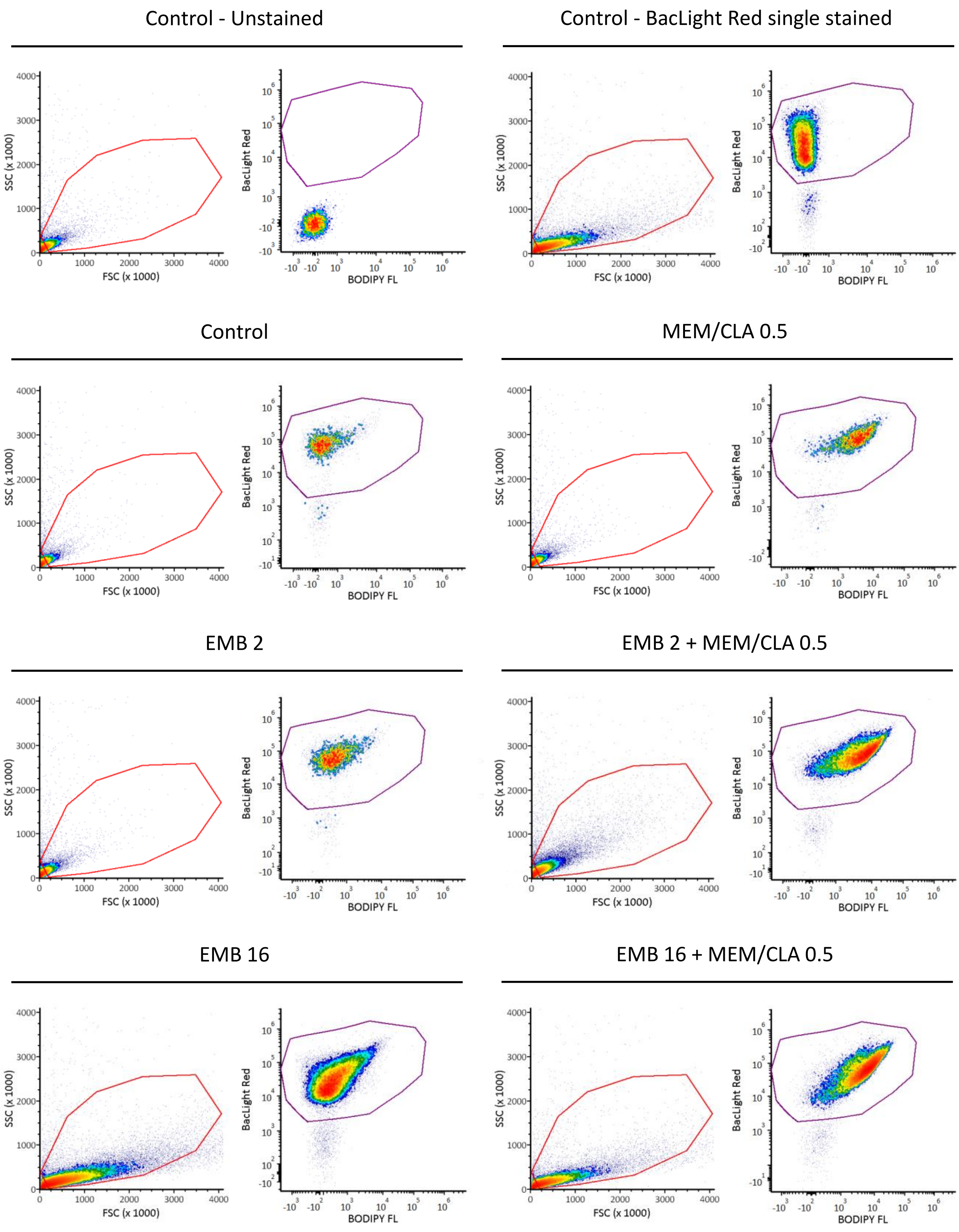


Figure S9. Representative replicates of the flow cytometry analysis of *Mycobacterium tuberculosis* strain MDR III present in Figures 5A-B. For each condition, the left panel shows the bacterial population gated for analysis (red gate) and the right panel shows the distribution of bacteria stained for BacLight Red and/or for BODIPY FL after 6 h of treatment. BODIPY FL fluorescence intensity was quantified from bacteria stained with BacLight Red (purple gate). CLA, clavulanate; EMB, ethambutol; MEM, meropenem. The numbers after each antibiotic abbreviature indicate the respective concentration in mg/L. When present, clavulanate concentration was fixed at 2.5 mg/L.

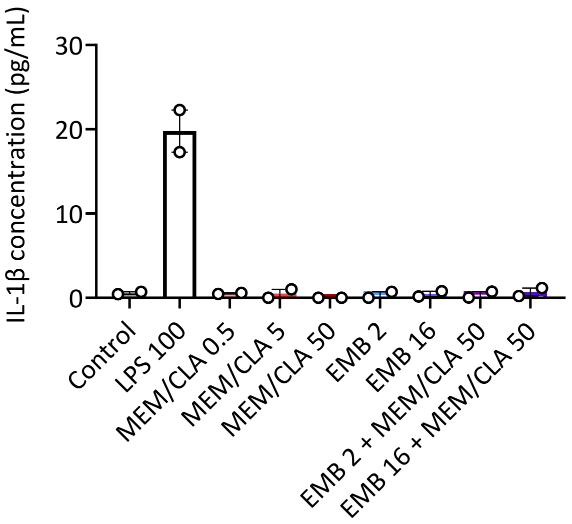


Figure S10. IL-1β quantification by ELISA in the supernatants of uninfected THP-1 macrophages after 24h of incubation with the antibiotics. Depicted values show the mean concentration of two replicates and error bars represent the standard error of the mean. CLA, clavulanate; EMB, ethambutol; MEM, meropenem. The numbers after each antibiotic abbreviature indicate the respective concentration in mg/L. When present, clavulanate concentration was fixed at 2.5 mg/L.

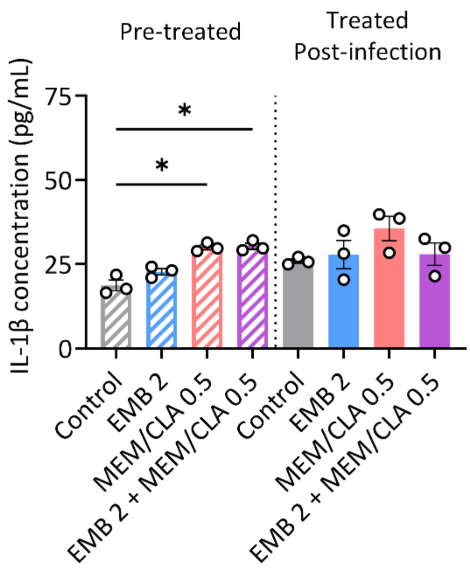


Figure S11. IL-1β quantification by ELISA in the supernatants of THP-1 macrophages infected with H37Rv pre-treated with the antibiotics for 6h ("Pre-treaded" samples) or with untreated H37Rv after 24 h of incubation with the antibiotics ("Treated Post-infection" samples). Depicted values show the mean concentration of three replicates and error bars represent the standard error of the mean. CLA, clavulanate; EMB, ethambutol; MEM, meropenem. The numbers after each antibiotic abbreviature indicate the respective concentration in mg/L. When present, clavulanate concentration was fixed at 2.5 mg/L.
